# Influenza A Virus H7 nanobody recognizes a conserved immunodominant epitope on hemagglutinin head and confers heterosubtypic protection

**DOI:** 10.1101/2024.10.31.621368

**Authors:** Zhao-Shan Chen, Hsiang-Chi Huang, Xiangkun Wang, Karin Schön, Yane Jia, Michael Lebens, Danica F. Besavilla, Janarthan R. Murti, Yanhong Ji, Aishe A. Sarshad, Guohua Deng, Qiyun Zhu, Davide Angeletti

## Abstract

Influenza remains a persistent global health challenge, largely due to the virus’ continuous antigenic drift and shift, which impede the development of a universal vaccine. To address this, the identification of broadly neutralizing antibodies and their epitopes is crucial. Nanobodies, with their unique characteristics and binding capacity, offer a promising avenue to identify such epitopes. Here, we isolated and purified a hemagglutinin (HA)-specific nanobody that recognizes an H7 subtype of influenza A virus. Notably, the nanobody, named E10, exhibited broad-spectrum binding, cross-group neutralization and *in vivo* protection across various influenza A subtypes. Through phage display and *in vitro* characterization, we demonstrated that E10 specifically targets an epitope on HA head. This epitope is part of the conserved lateral patch of HA head and proved to be highly immunodominant upon H7 infection. Importantly, immunization with a peptide including the E10 epitope elicited cross-reactive antibodies and mediated partial protection from lethal viral challenge. Our data highlight the potential of E10 and its associated epitope as a candidate for future influenza prevention strategies.

## Introduction

Influenza A (IAV) and B viruses are responsible for seasonal epidemics, leading to 290,000 to 650,000 human deaths globally each year<u>^1-3^</u>. Despite vaccination efforts, the continuous circulation of these viruses in humans, coupled with their ability to rapidly mutate, poses a significant ongoing public health threat. In addition, the current spread of H5N1 among various animal species, including cattle, is causing widespread concerns^4^. Similarly, avian influenza A (H7N9), circulating in birds and poultry, has led to over a thousand laboratory-confirmed human infections with a case-fatality rate of approximately 39% though human-to-human transmission has not been reported yet <u>^5,6^</u>. The H7N9 virus was first identified in China in March 2013<u>^7-9^</u>, with its the largest outbreak during the 5^th^ epidemic wave in 2016-2017<u>^10,11^</u> characterized by antigenic drift in the hemagglutinin (HA) protein. H7N9 may even possess greater pandemic potential than H5N1, highlighting the urgency of developing effective therapeutic strategies <u>^12^</u>. Currently, the antiviral treatment for H7N9 are limited to neuraminidase (NA) inhibitors <u>^13^</u>. Vaccination remains the most effective way of preventing IAV<u>^14^</u> reducing the risk of illness by 40% to 60%<u>^15,16^</u>. However, it is crucial to identify conserved sites of vulnerability in both human and avian influenza viruses to develop antibody therapy or novel vaccines.

Influenza HA is the immunodominant surface glycoprotein of IAV and targets of most of the antibody (Ab) response. HA can be broadly divided into globular head and stem domains<u>^17^</u>. Most of the Ab response is targeting variable regions such as the canonical antigenic sites in HA head <u>^18-21^</u>. On the other hand, stem-specific Abs exhibit higher breadth and are often capable of neutralizing several IAV strains<u>^22,23^</u>. Indeed, cross-neutralizing stem Abs can provide a further line of defense against the virus<u>^17,24,25^</u>. Of note, while Abs to head are mostly strain specific, several broadly neutralizing HA-head Abs have also been identified<u>^26-28^</u>, underscoring the importance of targeting conserved epitopes in the development of universal vaccines. Among those, Abs targeting the receptor-binding site (RBS) and the lateral patch on H1<u>^29,30^</u>. Yet, there is limited knowledge about lateral patch-binding Abs upon H7N9 infection<u>^5^</u>. In general, the characterization of the epitopes of broadly neutralizing antibodies has greatly aided the development of several universal influenza vaccine candidates by selecting conserved epitopes<u>^28,31,32^</u>.

Beside Abs, nanobodies<u>^33^</u>, derived from camelids, offer a promising alternative due to the small size (15 kDa)<u>^34^</u>, strong physical<u>^35^</u> and chemical stability<u>^36^</u>, and ability to access cryptic viral sites and enhance tissue permeability<u>^37-40^</u>. Furthermore, nanobodies are easier to produce and are poorly immunogenic in humans<u>^41,42^</u>. A broadly neutralizing nanobody targeting multiple influenza subtypes could be a valuable addition to our therapeutic arsenal, particularly in the event of a new pandemic<u>^43-45^</u>.

However, immunodominance in B cell responses often steers Ab away from these conserved regions<u>^46^</u>, directing them to variable areas of HA head instead. Therefore, understanding the mechanisms of immunodominance is essential for redirecting Abs responses to desired targets within HA<u>^47^</u>. While HA immunodominance in H1N1 and H3N2 is well-characterized<u>^18,48,49^</u>, it remains largely unexplored in H7N9, especially regarding how antigenic drift redirects B cell and Ab response to H7N9<u>^50,51^</u>. Indeed, absence of strongly immunodominant sites may be beneficial in allowing the Ab response to more equally spread across several targets.

In our study, starting with H7 HA immunization, we identified a broadly neutralizing and broadly protective nanobody targeting the lateral patch region of HA. Furthermore, immunization with the corresponding epitope elicited a cross-reactive, protective response. Our results provide a potential therapeutic tool and a blueprint for designing an effective universal vaccine against influenza viruses with pandemic potential.

## Results

### Generation and characterization of H7-specific nanobodies

The H7N9 virus is a rapidly evolving virus with similar pandemic potential as H5N1<u>^5,52^</u>, making it a particularly attractive, but challenging, target to study. Indeed, previous studies on nanobodies have primarily focused on H1N1 and H3N2 HA proteins<u>^25,30^</u>. We selected H7N9 virus A/Environment/Suzhou/SZ19/2014 (SZ19)<u>^53^</u>, previously isolated in our lab, as a well-characterized strain with high relevance to recent outbreaks, making it an ideal candidate for studying the effectiveness of H7-specific nanobodies.

To identify SZ19 H7-specific nanobodies, we immunized alpacas five times with H7N9 virus (Fig. 1A). After 14 days from the last immunization, we collected peripheral blood mononuclear cells (PBMCs) and constructed a phage display library. Using SZ19 H7-HA protein as the target, we performed bio-panning three rounds, leading to selection of 96 colonies. Among them, we sequenced the top 20 binders and identified six unique nanobodies (A11, C11, E10, F3, H10, and H12), all expressed with Fc tags for enhanced functionality. (Figs. 1A and Fig S1A, highlighted in yellow). As shown by the molecular model<u>^54^</u> nanobodies have one single-domain, and even after linking with Fc tag the total size is only about 40 kilodaltons (kDa) (in Fig. 1B and Fig. S1B-C), which is remarkably smaller than a regular Ab (150 kDa)<u>^55^</u>. While F3 and H10 appeared closely related, all other nanobodies, presumably came from different precursors; E10 stood out as it had the longest CDR3 and for the presence of several negatively charged amino acids (Fig. S1F). To test their ability to recognize folded viral HA, we performed immunofluorescence assay (IFA) as well as western blotting (WB) at different time points after infection (Figs. 1C-D, S1D-E). All six nanobodies were able to recognize HA when linearized in WB (Figs. 1D, S1E) and the cytoplasm of SZ19 virus-infected cells (Fig. 1C), suggesting that they all target a linear epitope which is also present on nascent HA.

**Figure 1.**
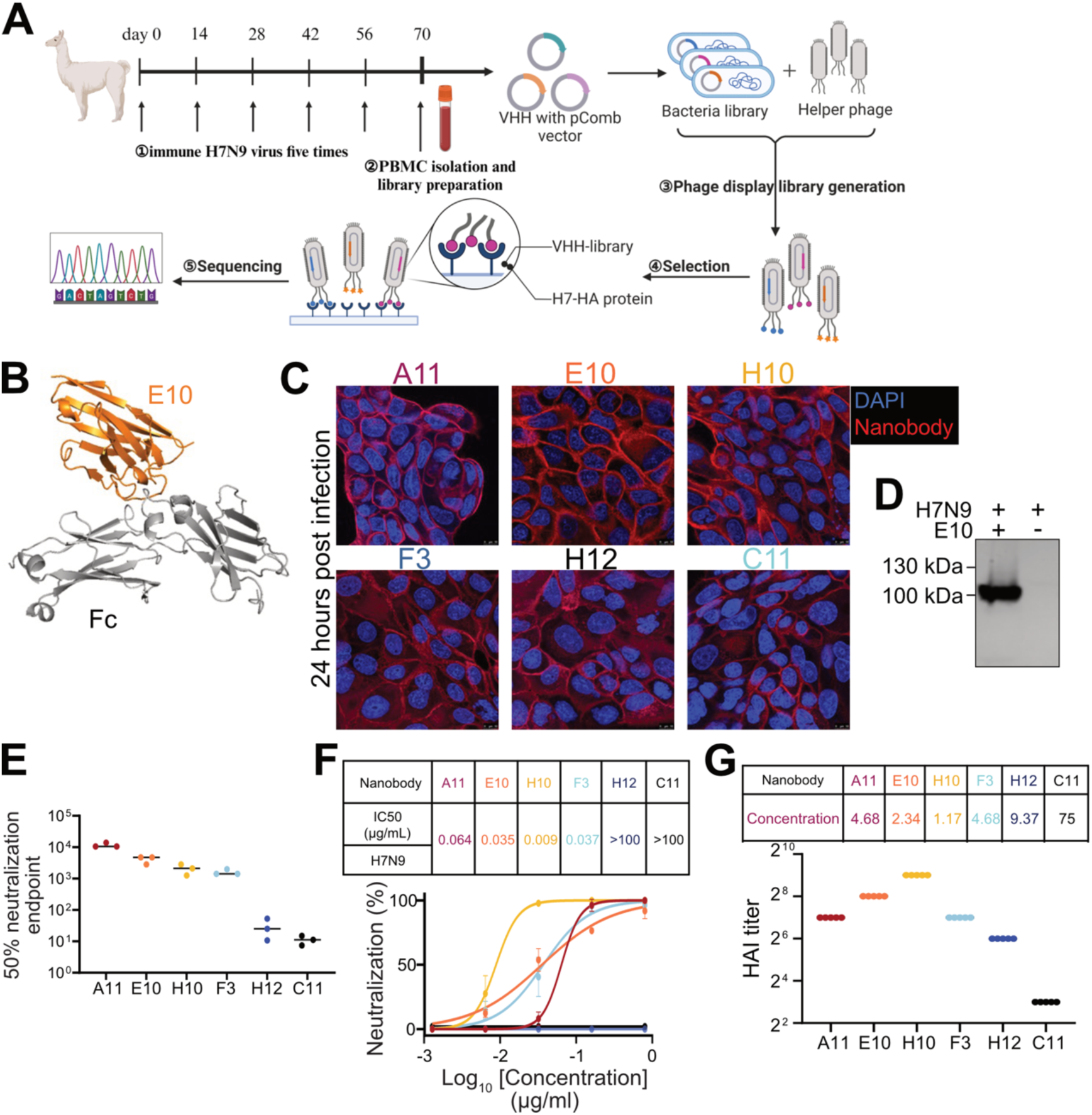
Generation and characterization of H7-specific nanobodies. (A) Schematic illustration of alpaca immunization process in alpacas and subsequent nanobody selection. Alpacas were immunized with the H7N9 virus, and after five rounds, peripheral blood mononuclear cells (PBMCs) were collected to generate a phage display library. (B) Structural model of a nanobody-Fc created using ImmuneBuilder and AlphaFold 2. (Nanobody: orange one, we select E10 to show the model. Fc tags: gray.). (Nanobody: orange one, we select E10 to show the model. Fc tags: gray.) (C) Immunofluorescence assay (IFA) showing the recognition of the SZ19 virus by different nanobodies. A549 cells were infected with SZ19 (MOI=0.1) for 24h and stained with distinct primary nanobodies, followed by secondary goat anti-human IgG Fc Alexa Fluor™ 488 (A11: red, E10: orange, H10: yellow, F3: blue, H12: dark blue, C11: black.). (D) Western blot (WB) analysis of A549 cells infected with SZ19 (MOI=1) for 24h. Cell lysate was probed with respective nanobodies and detected using goat anti-human IgG-Fc HRP. Data is representative of at least two independent experiments. (E) Graph showing 50% neutralization endpoint (NC_50_) of different nanobodies against the SZ19 as measured by their ability to inhibit hemagglutination of red blood cells (RBCs). Data is representative of at least two independent experiments. Shown are the mean values of three replicates. (F) Neutralization assay results for the nanobodies on SZ19 using a cell-based ELISA. Half-maximal inhibitory concentrations (IC_50_) are shown for each nanobody. Data represents the mean values from four technical replicates in at least two independent experiments. (G) Hemagglutination inhibition assay (HAI) tests the binding area of six different nanobodies in H7N9-HA protein (A11: red, E10: orange, H10: yellow, F3: blue, H12: dark blue, C11: black. All the nanobodies colors are the same as blow.). We diluted nanobodies two-fold starting from 300ng/µL. Data is the average of three independent experiments.

Neutralizing Abs can block viral spread by preventing infection<u>^56^</u>, and it is indeed desirable for nanobodies to be neutralizing<u>^57^</u>. To test neutralization, we incubated various concentrations of nanobodies with H7N9 viruses at 37°C for one hour (1h) before adding them to Madin-Darby Canine Kidney (MDCK) cells. Thereafter, we measured viral replication using two methods. First, we took the supernatant after 72h and measured 50% neutralization titer (NC_50_) using hemagglutination assay (Fig. 1E). Alternatively, we waited for 20h and subjected infected MDCK to cell-based ELISA using an NP-specific Ab (Fig. 1F). Both methods showed similar results, indicating that four nanobodies were neutralizing (A11, E10, H10, F3), while H12 and C11 were not. Calculation of 50% inhibitory concentration (IC_50_) by cell-based ELISA demonstrated that A11, E10 and F3 had similar potency, while H10 had an approximately 4-7-fold better neutralization potency (Fig. 1F).

To further explore the characteristics of nanobodies, we sought to map out the nanobodies binding site and see whether they bind close to HA-head or stalk. As a first rough indication we used hemagglutination inhibition test<u>^58^</u>. All neutralizing nanobodies (A11, E10, H10, F3) were able to inhibit hemagglutination (Fig. 1G) with a high HI titer. Surprisingly, even the non-neutralizing nanobody H12 demonstrated HI activity. Overall, the results suggested that all nanobodies, except C11, bind to HA-head.

### Nanobody E10 exhibits cross-group binding and neutralization capacity

Following the identification of four potent nanobodies capable of neutralizing the homologous H7N9 virus, we expanded our analysis to determine if any of these could also recognize and neutralize other IAV strains. HA is phylogenetically divided into two major groups: group one, including circulating H1, and group two, comprising circulating H3 and avian H7 strains<u>^59^</u>. Typically, antibodies targeting the HA head are strain-specific<u>^60^</u>, however, some can recognize conserved epitopes across multiple subtypes, offering cross-group neutralization<u>^61^</u>. Nanobodies, due to their unique prolate (rugby ball-shaped) structure and compact variable heavy-chain domain(VHH), form a convex paratope surface<u>^33^</u>, with higher possibility to access antigen cavities that are often hidden from conventional antibodies, increasing their potential to recognize conserved epitopes invisible to human Abs.

First, to evaluate the binding capacity of the six nanobodies, we performed ELISA on plates coated either with either UV-inactivated virus or recombinant HA proteins. We tested not only H7 HA but also A/Puerto Rico/8/1934 (PR8, H1N1) H1 virus and A/Hong Kong/1968 (X31, H3N2) H3 viruses and their respective HA proteins. The nanobodies displayed varying binding ability across different viral subtypes and HA proteins. Importantly, nanobody E10 consistently demonstrated strong binding to all viruses and HA proteins tested, suggesting broad binding capacity (Figs. 2A-F).

**Figure 2.**
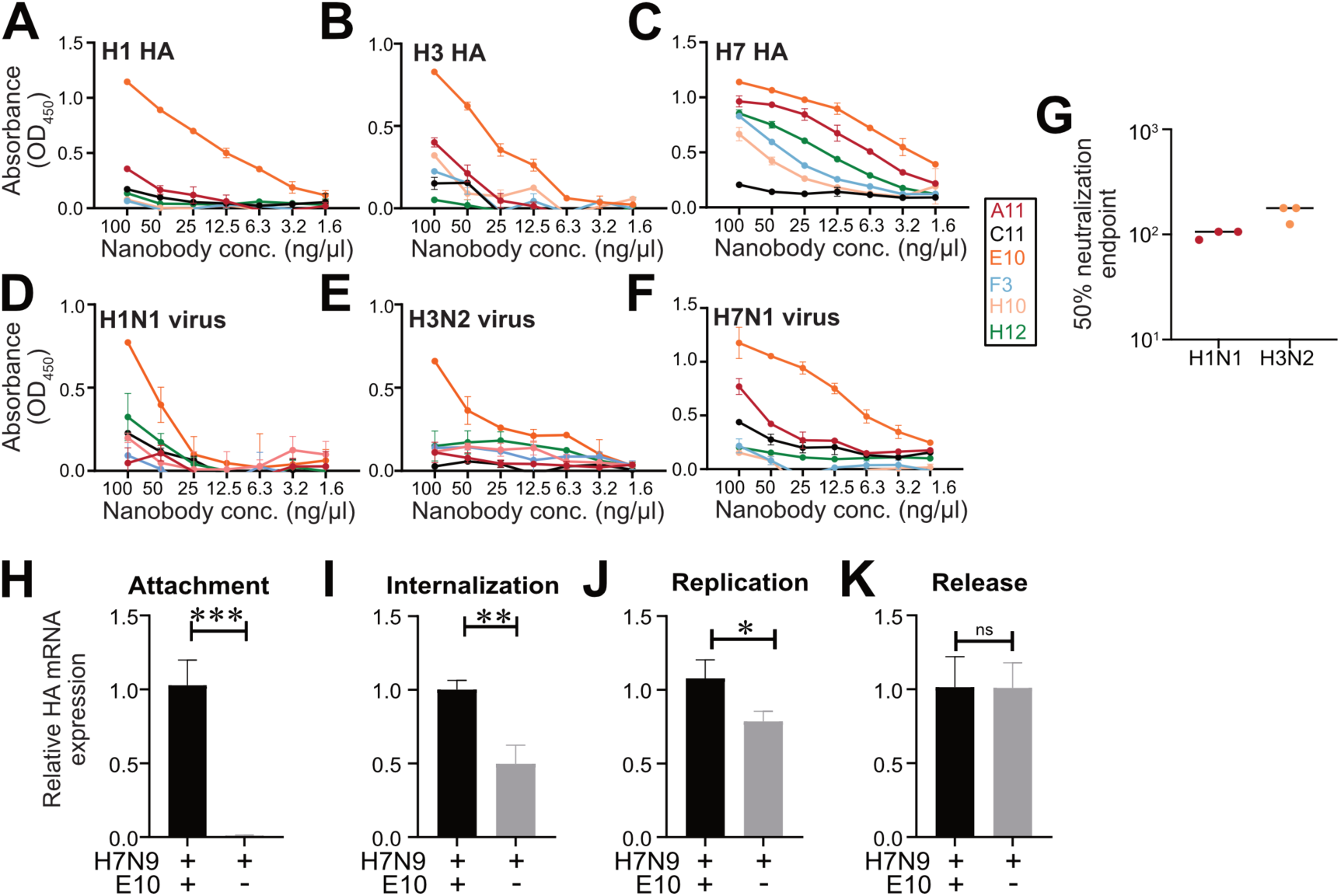
Nanobody E10 exhibits cross-group binding breadth and neutralization capacity. (A-F) ELISA binding curves showing the interaction of six nanobodies with HA protein from various influenza strains (H7, H1, and H3) and UV-inactivated influenza viruses (H7N1, H1N1, and H3N2). Each curve represents a different nanobody, color-coded for clarity. Data are presented as mean ± SD from at least two independent experiments. (G) Neutralization efficacy of nanobody E10 against different subtypes of influenza A virus (H1N1 and H3N2). Data are representative of at least two independent experiments. Shown are the mean values of three technical replicates. (H-K) Evaluation of E10’s effect on different stages of the viral infection process using SZ19 (MOI=10) at various time points. Bars represent the mean HA gene expression levels measured by qPCR. Data are shown as mean values from three technical replicates. Statistical significance was determined using an unpaired t-test: *p < 0.05; **p < 0.01; ****p < 0.0001; ns: not significant.

We further examined the neutralization capacity of E10 using the methodology described for Fig. 1E, and it demonstrated effective neutralization of H1N1 and H3N2 (Fig. 2G). These results highlighted E10 outperformed other nanobodies, demonstrating remarkable cross-group binding and neutralization capabilities. Given these promising findings, the rest of the study focused primarily on unraveling the mechanisms behind E10’s broad efficacy.

The life cycle of influenza virus can be divided into four key stages: attachment, internalization, genome replication, and viral release. Most antibodies targeting the HA head prevent viral entry by blocking attachment<u>^56^</u>. To determine whether E10 acted via a similar mechanism, we first tested its ability to block viral attachment. We pre-incubated E10 with SZ19 at 37℃ for 1 hour, then added the mixture to A549 cells at 4℃ for 1hour to allow binding without internalization (Fig. 2H). The results clearly showed that E10 efficiently blocked viral attachment. Next, to verify E10 ability to block viral internalization and gene replication, we followed the same protocol but added a 4h (internalization) or 8h (gene replication) incubation at 37℃, after adding it to cells (Figs. 2I-J). Finally, to check whether E10 could inhibit viral release, we added the nanobody after allowing viral infection for 10 hours (Fig 2K). The most pronounced effect of E10 was on attachment, with some effect on viral internalization but no influence on viral release, thus establishing its function in inhibiting attachment and entry.

### E10 treatment protects mice against homo- and heterosubtypic IAV challenge

Following the *in vitro* demonstration of E10’s capacity to neutralize multiple influenza strains by blocking viral attachment to host cells, we proceeded to evaluate its efficacy *in vivo*. For prophylactic assessment, we administered 10 mg/kg of E10 to mice for 24 h, followed by infection with 10^6^ EID_50_ of SZ19 H7N9 virus (Fig. 3A). In parallel, we assessed the therapeutic potential by administering 25 mg/kg of E10 2h post IAV infection (Fig. 3B). Mice were either sacrificed on day 3 to check for lungs pathology and viral titers in various organs, or their weight was monitored over 14 days, with a 25% weight loss used as a cutoff.

**Figure 3.**
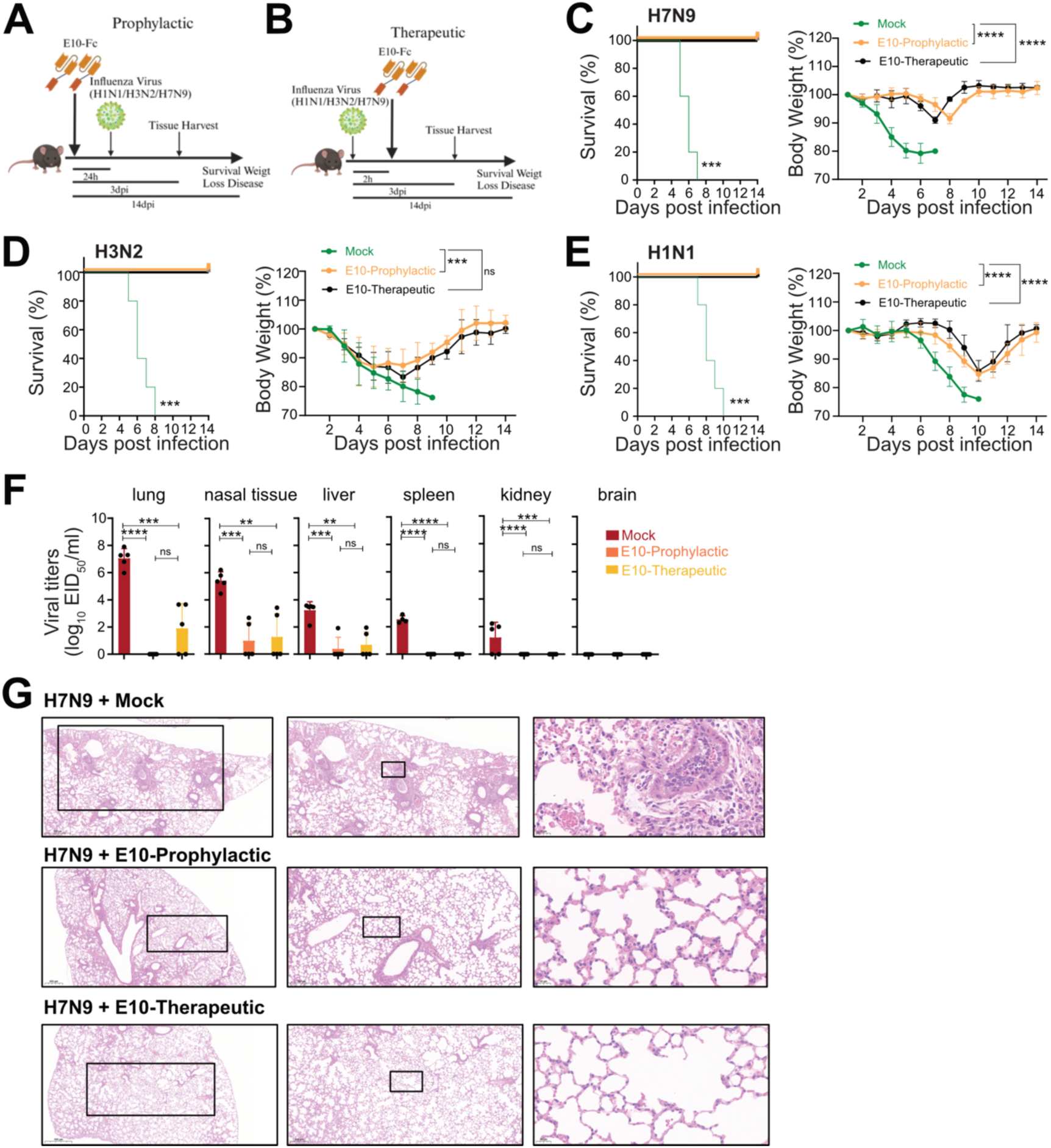
E10 treatment protects mice against homo- and heterosubtypic IAV challenge. (A-B) Schematic representation of the experimental design for *in vivo* studies. Mice were treated with either E10-Fc or PBS intraperitoneally (i.p.), followed by influenza virus infection (H1N1, H3N2 or H7N9), as outlined. (C-E) Kaplan-Meier survival curve and body weight monitoring of influenza-infected mice. Mice were treated with E10-Fc or PBS as prophylaxis or therapy i.p and weight monitored for 14 days after infection with H1N1 (3×10^3^ TCID_50,_ lethal dose)/ H3N2 (10^7^ TCID_50,_ lethal dose) / or H7N9 (10^6^ EID_50,_ lethal dose). Statistical significance of the Kaplan-Meier survival curves was calculated by log rank Mantel-Cox test. ∗p < 0.05, ∗∗p < 0.01, ∗∗∗p < 0.001, ∗∗∗∗p < 0.0001, ns: not significant. Weight change data are represented as mean ± SEM from three independent experiments (n = 15 per group). Statistical analysis was performed using two-way ANOVA: ∗p < 0.05, ∗∗p < 0.01, ∗∗∗p < 0.001, ∗∗∗∗p < 0.0001, ns: not significant. (F) Viral titer, measured by EID_50_, in six organs (lungs, nasal tissue, liver, spleen, kidney, brain) on day 3 post-H7N9 infection, with or without E10 treatment, as described in A-B. Data are representative of three independent experiments and shown as mean values from three technical replicates. Statistical analysis was performed using unpaired t test. *p < 0.05; **p < 0.01; ****p < 0.0001; ns: not significant. (G) Representative histopathological analysis of lungs from mice on day 3 after H7N9 infection with or without E10, and with MOCK mice, as outlined in A-B. Scale bar, 500 μm (left), 200 μm (middle), 20 μm (right).

E10 treatment significantly prevented weight loss and mortality, both when administered before and after infection (Fig 3C). In untreated mice, H7N9 spread to multiple organs, including the liver, spleen, and kidneys (Fig 3F). However, E10 treatment completely abolished viral replication in most organs, including the lungs, with only a few mice showing viral titers in nasal tissues, indicating that E10 effectively protects various organs *in vivo* (Fig. 3F). As expected, therapeutic administration did not entirely prevent viral replication in the lungs, but it did reduce viral loads by 5 logs compared to untreated mice (Fig. 3F). Lung pathology analysis further confirmed the protective effect of E10, regardless of timing of administration. While H7N9 infected, untreated mice exhibited obvious lung lesions and granulocyte infiltration by day 3 post-infection, mice treated with E10, either prophylactically or therapeutically, showed normal lung tissue architecture with no visible signs of immune cell infiltration (Fig. 3G)

Given E10’s cross-neutralization ability *in vitro,* we tested heterosubtypic protection *in vivo*. We used mouse adapted versions of two IAV subtypes currently circulating in humans, belonging to two distinct phylogenetic groups. Animals were infected with either X31 (H3N2) or PR8 (H1N1), with E10 administered before or after infection and their weights monitored (Fig 3D-E). All untreated, infected mice succumbed to infection by day 9 (H3N2) or day 10 (H1N1). However, consistent with its *in vitro* activity, E10 was able to convey protection against both strains, reducing mortality to zero, despite some weight loss, even in animals which received the nanobody (Fig 3E-J). Likewise, E10 administration also significantly reduced viral titers and lung pathology in heterosubtypic infections (Figs. S2A-B).

### E10 recognizes a conserved epitope located on HA-head lateral patch

To identify the binding site of nanobodies, we generated escape virus variants using 10- days-old specific pathogen-free (SPF) eggs (Figs. 4A, S3A). This approach takes advantage of the error-prone polymerase of IAV to determine the Ab-epitopes<u>^62^</u>. Different dilutions of each nanobody were mixed with H7N9 virus and injected into the eggs. After three to four rounds of selection, hemagglutination inhibition-positive supernatants were submitted for next-generation sequencing (NGS). E10-selected escape mutants at residues K166T and S167L (H3 numbering, used throughout the manuscript), A11-selected escapes at K140N, while H10 at residues S167L and G205E.

**Figure 4.**
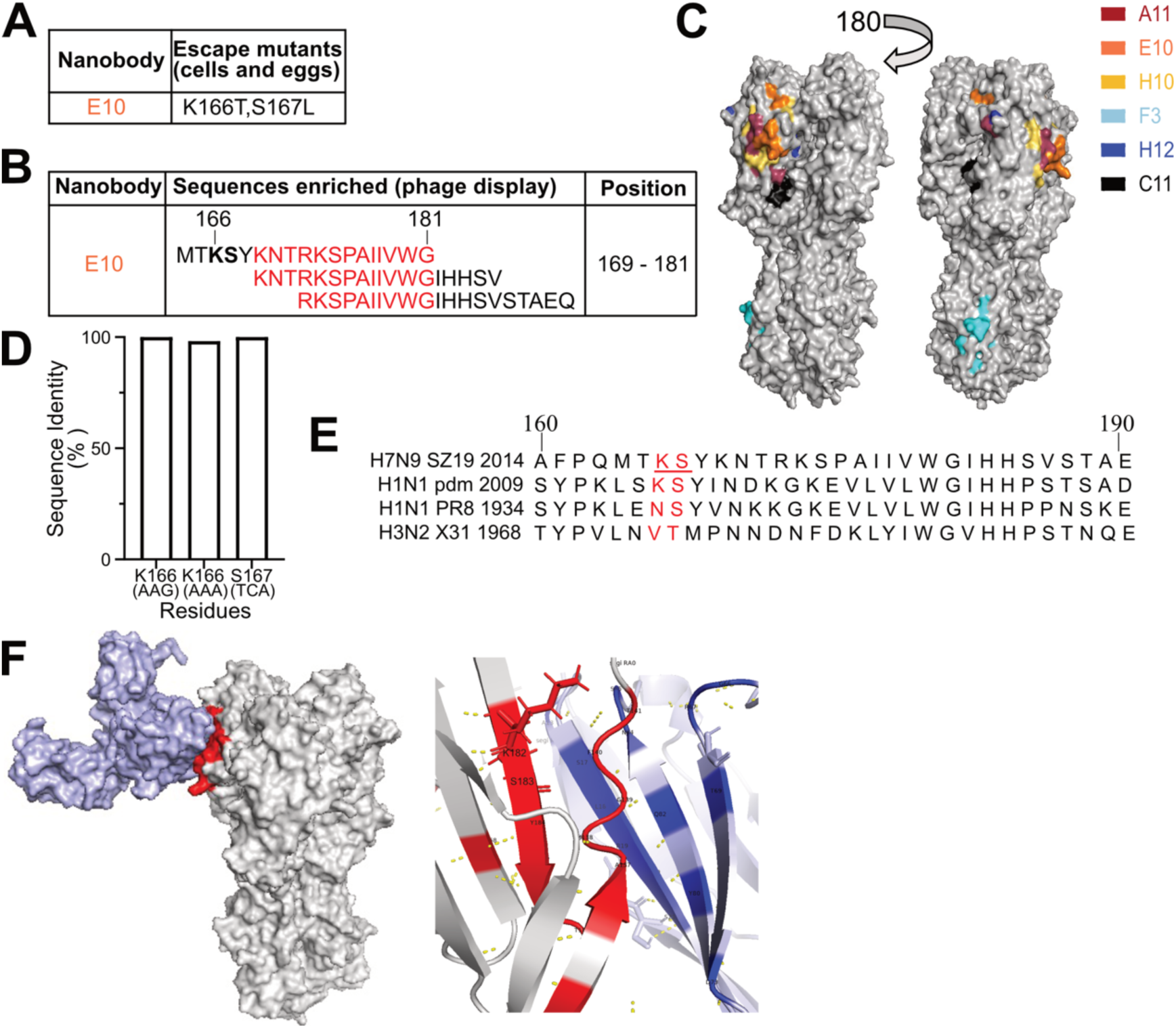
E10 recognizes a conserved epitope located on HA-head lateral patch. (A) E10 escape mutations in the H7N9 virus were identified using SPF eggs and MDCK cells. Variations at key residues involved in nanobody escape are highlighted, showing the sites where resistance developed. (B) Phage display selection of E10 nanobody identified specific peptides. The red region depicts an overlapping area among selected peptides. The bolded region corresponds to the epitope identified through escape mutations, which overlap with key binding residues. (C) Epitope mapping of the H7N9 HA protein was performed, showing the binding sites of six different nanobodies, each labeled in distinct colors (A11: red, E10: orange, H10: yellow, F3: blue, H12: dark blue, C11: black). The head domain of H7 HA was modeled using Swiss-Model. Images were generated using PyMOL software. (D) Quantitative analysis of amino acid substitutions at positions 166 and 167 across all H7N9 strains in the GISAID database. (E) Amino acid sequences flanking the E10 epitope are compared across various IAV strains, including A/Environment/Suzhou/SZ19/2014 (H7N9), A/California/07/2009 (H1N1 Pandemic), A/Puerto Rico/8/1934 (PR8, H1N1), and A/Hong Kong/1968 (X31, H3N2). Variations at positions 166 and 167 are highlighted in red, with the E10 epitope underlined. SZ19 HA derivatives with the corresponding substitutions are also included for comparison. (F) Three-dimensional (3D) structure analysis of intramolecular interactions between E10-Fc and H7N9-HA. The red area depicts the amino acid interactions of H7-HA between E10-Fc and H7-HA, K182 and S183 sticks (K166 and S167 in H3 numbering) marked in H7N9-HA is the main binding sites selected by eggs/cells. The blue area depicts the amino acid interactions of E10-Fc between E10-Fc and H7-HA. Purple represents E10-Fc, and gray represents the HA protein.

Furthermore, we repeated the selection process for E10 using cell culture, and obtained the same escape mutants, thus confirming its binding site (Figs. 4A, S3A). Cell culture escape selection results for A11 and H10 were comparable to those obtained in eggs, with H10 additionally selecting for a mutation at residue S167, while H12 now selected for mutants. Furthermore, we constructed an H7 fragment library for phage display selection, utilizing 20-mer peptides overlapping by 5 amino acids. We panned phages using all nanobodies (Fig S3B) and identified binding sites which were in agreement with those selected in cells and egg. Of note, E10 selected for three overlapping peptides, one of them including the K166 and S167 residues (Fig. 4B) and the others in close proximity. Altogether, by combining the three different methods we mapped all the nanobodies epitopes on a molecular model of H7 HA (Figs. 4C, S3C-D). Interestingly, E10, A11 and H10 bound in a similar region near the lateral patch of HA while the F3 epitope localized to the stem of HA.

Conservation analysis of the E10 mutated residues showed that both K166 and S167 are highly conserved across H7-HA proteins (Fig. 4D), suggesting limited immune pressure on this site or lower viral fitness of escape mutants. While these residues were fully conserved in the currently circulating pandemic H1N1, some variation was observed within H1N1-HA (PR8) and H3N2-HA (X31) (Fig. 4E). However, the ∼20 amino acids close to that region showed a good degree of conservation, suggesting that the nanobody footprint may also include nearby residues, as also suggested by the phage display selection. To verify this, we modeled the interaction between E10-Fc and H7-HA: indeed most of the contact surface of HA, with the nanobody included conserved β-sheet structures on the lateral patch of HA (Fig. 4F). In summary, our combined mutation and epitope mapping analyses demonstrated that the broadly reactive E10 nanobody recognizes a highly conserved epitope on the lateral patch of H7 HA.

### H7-HA_K166T, S167L_ mutant virus has reduced viral fitness

To confirm that the selected escape mutations were indeed part of the E10 binding site, we employed multiple experimental approaches. First, we verified E10-Fc recognition on H7-infected A549 cells and while E10 readily stained H7N9-infected cells (thereafter referred as WT virus), we could not detect any binding in H7N9_K166T, S167L_-infected cells (referred to as MUT virus) (Fig. 5A). Likewise, escape mutants obtained by A11 selection failed to recognize their respective mutated virus (H7HA_K140N_) (Fig. S4A). Since E10 recognizes a linear epitope, we further verified the escape of MUT virus by WB. Here, we incubated WT or MUT virus with E10 prior to infecting cells and detected productive infection by NP antibody and HA recognition by E10. WT virus without E10 pre-incubation readily infected cells and HA was detected by the nanobody. Notably, pre-incubation with E10 readily blocked infectivity of WT but not MUT virus and E10-nanobody was not able to react with MUT-HA (Fig 5B, S4B, S4C). Further confirming escape, E10 was not able to neutralize MUT virus infection *in vitro* (Fig 5C). Finally, we expressed H7 HA from WT and MUT viruses as recombinant HA proteins (thereafter referred to as WT-HA protein and MUT-HA protein) (Fig S4D) and tested nanobody reactivity. As expected, E10 lost all reactivity towards MUT-HA while F3 control nanobody recognized both proteins equally (Fig 5D).

**Figure 5.**
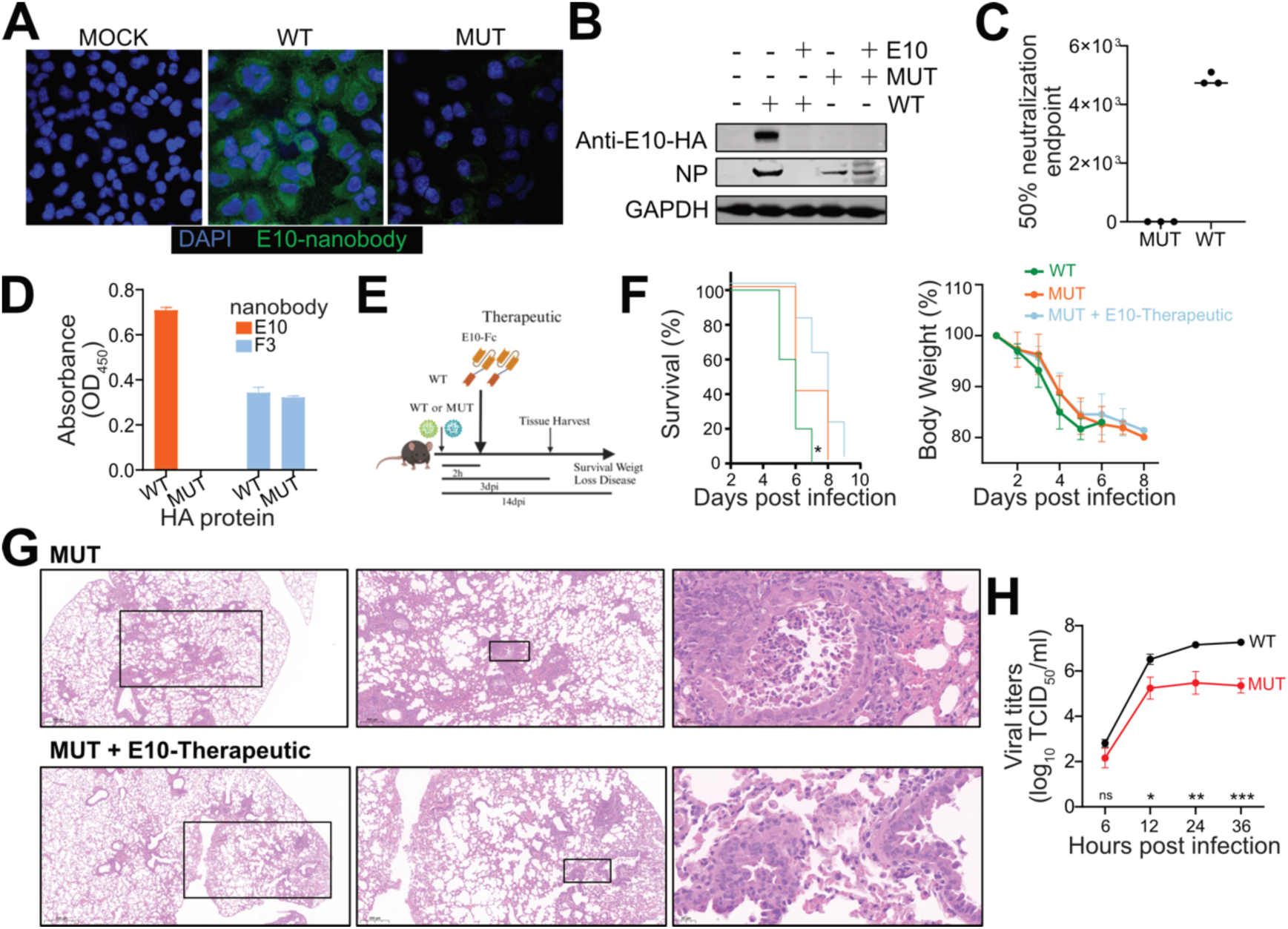
H7HA_K166T, S167L_ double mutant escapes E10 recognition but has lower viral fitness. (A) IFA of A549 infected with wild-type (WT) or mutant (MUT) H7N9 virus (MOI = 0.1) at 24 hours post-infection. Cell was stained with E10 as the primary antibody, followed by secondary goat anti-human IgG Fc Alexa Fluor™ 488. Nuclei were counterstained with DAPI (blue), and NP nanobody detection is shown in green. Data is representative of at least two independent experiments. (B) WB analysis of A549 cells infected with WT or MUT H7N9 viruses, with or without E10 pre-incubation, showing detection of NP and E10. Data is representative of at least two independent experiments. (C) Graph showing the 50% neutralization endpoint of E10 against WT and MUT viruses, measured by the ability of virus progeny to hemagglutinate red blood cells (RBCs). Data is representative of at least two independent experiments. Shown are the mean values of three technical replicates. (D) ELISA detection of WT-HA and MUT-HA hemagglutinin (HA) by E10 and F3 nanobodies. Bars represent mean ± SD. Data is representative of at least two independent experiments. Shown are the mean values of three technical replicates. (E) Experimental setup for the studies shown in panels F and G. (F) Kaplan-Meier survival curve and body weight analysis of influenza-infected animals. Mice were treated intraperitoneally (i.p.) with either E10-Fc or PBS, as depicted in panel E, and their weight was monitored for 14 days following infection with WT (10⁶ EID_50_, lethal dose) or MUT virus (10⁶ EID_50_). The statistical significance was calculated by log rank Mantel-Cox test for survival curve, ∗p < 0.05, ∗∗p < 0.01, ∗∗∗p < 0.001, ∗∗∗∗p < 0.0001, ns (not significant). Weight change was monitored, and each graph is three experiments; n = 15; two-way ANOVA: ***P = 0.0002; ****P < 0.0001; symbols represent means ± SEM) (G) Representative histopathological analysis of mouse lungs at day 3 post-infection with MUT virus, following E10 administration as outlined in 5E. Scar bar, 500 μm in the left row, 200 μm in the middle row, 20 μm in the right row. (H) Growth kinetics of WT and MUT viruses in MDCK cells. Supernatants were collected at 6-, 12-, 24-, and 36-hours post-infection (MOI = 0.001). Virus titers are presented as mean ± SD from three independent experiments. \**P* < 0.01 (two-way ANOVA followed by Dunnett’s test).

Finally, to test the *in vivo* ability of E10 to protect against MUT virus we conducted the same experiments as in Fig 3 and determined its prophylactic and therapeutic activity upon lethal viral challenge (Fig 5E). Again, mice were either sacrificed on day 3 to check for lung pathology and viral titers in different organs, or their weight monitored for 14 days (Fig 5F, 5G and Fig S4E). The *in vivo* results confirmed the loss of efficacy of E10 on the MUT virus, cautioning that an escape to this broadly neutralizing nanobody is possible. However, MUT showed a slightly decreased pathogenicity *in vivo*, including a longer survival and lower lung viral titer, when compared to WT. These results suggest that MUT may have a less virulent phenotype, thus raising hopes for the targeting of this epitope. To confirm this, we compared the *in vitro* growth kinetics of WT and MUT viruses: WT grew rapidly and reached peak titer by 36h post infection where it killed out most cells. Conversely MUT virus grew slower and to a peak over two logs lower than WT (Fig 5H), demonstrating a reduced viral fitness. Overall, while the emergence of escape mutants in response to E10 nanobody is possible, their viral fitness was reduced both *in vitro* and *in vivo*.

### E10-epitope is immunodominant upon H7-IAV infection

Little is known about H7 B cell immunodominance. Previous research suggested that single amino acid differences may alter the establishment of broadly neutralizing B cells<u>^63^</u>. Additionally, studies on immunodominance in humans have shown that antibody responses can be highly focused on specific epitopes, and that a single amino acid mutation can allow the virus to escape immunity in some individuals<u>^64-67^</u>. We reasoned that with our current tools we would be able to dissect the immunodominance of the B cell responses to the E10 epitope upon infection.

To this end we infected mice with either WT or MUT virus and we carried out B cell staining of lungs, the draining mediastinal lymph node (mln) and the spleen by flow cytometry <u>^68^</u>. Beside classical markers to distinguish germinal center (GC) and memory (MBC) B cells, we also included a multiple HA staining to define epitope specificity (Fig 6A, 6C, S5A). All cells were stained with WT-HA labelled with two distinct fluorophores and MUT-HA with other two fluorophores: for WT infected mice, we gated first on WT-HA double positive, as this was the only protein seen by the animals. Thereafter we used the staining with the MUT-HA to discriminate how many B cells were specific for the E10 epitope (Fig 6B, 6D): indeed, if WT-HA positive cells lost the recognition when stained with MUT-HA this suggested a specificity for the E10 epitope. Likewise, for mice infected with MUT virus, we first gated on MUT-HA and thereafter only the ones not recognizing the WT-HA were defined as specific for the mutated E10 epitope (Fig 6B, 6D).

**Figure 6.**
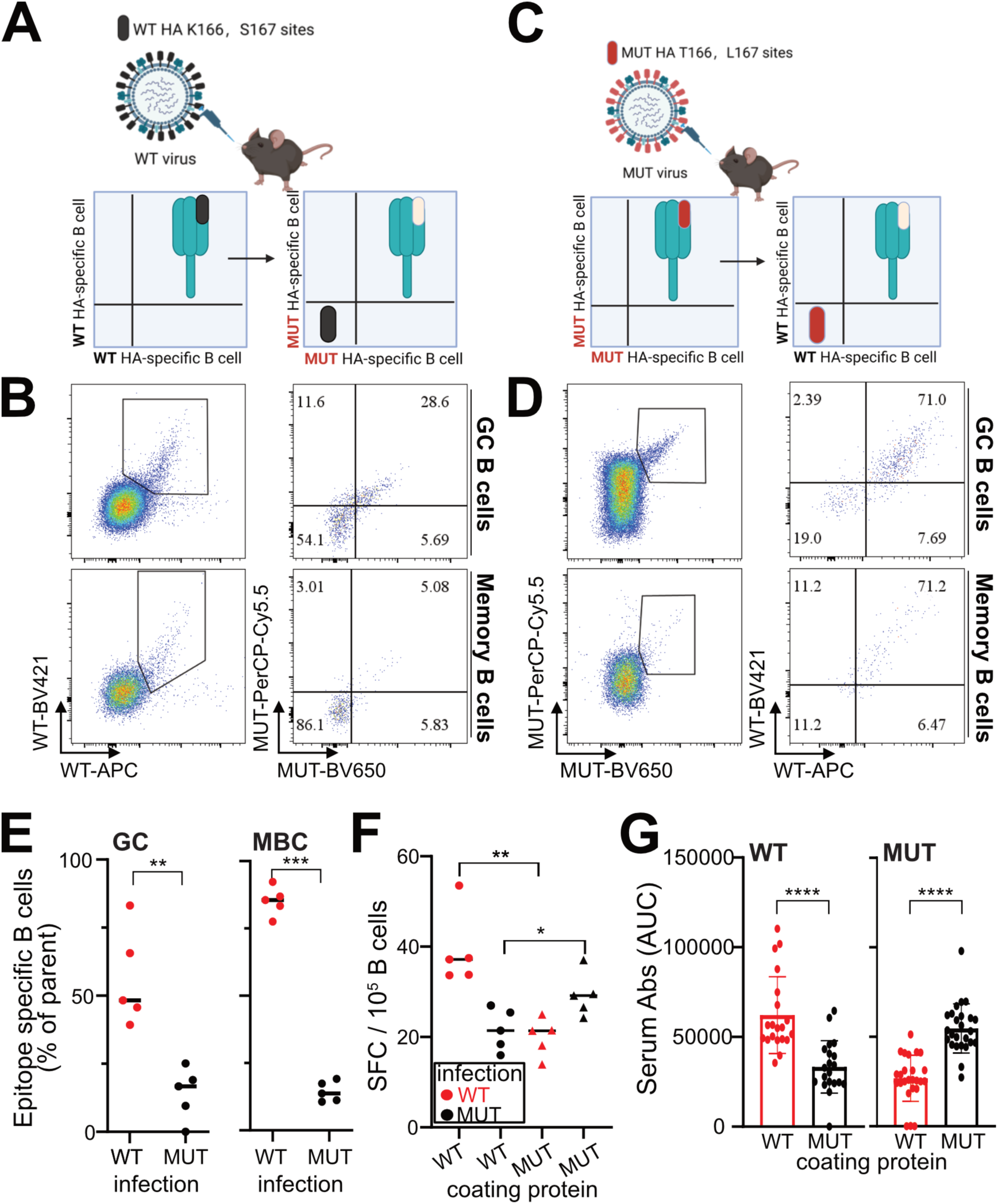
E10-epitope in immunodominant upon H7-IAV infection. (A, C) Schematic illustration of epitope identification by flow cytometry. (B) Representative flow cytometry gating of GC and MBC for epitope identification in mln of WT-infected mice 14 days post-infection. GC B cells were gated as live CD3− B220+ IgD− IgM− GL7+ CD38− WT+ MUT+, MBC as live CD3− B220+ IgD− IgM− GL7− CD38+ WT+ MUT+. (D) Same as in panel B but for MUT-infected mice, with GC B cells gated as live CD3− B220+ IgD− IgM− GL7+ CD38− MUT+ WT+, and MBC as live CD3−B220+ IgD− IgM− GL7− CD38+ MUT+ WT+. (E) Quantification of epitope-specific MBC and GC B cells in mln of WT- and MUT-infected mice at 14 days post-infection. Two independent experiments with 5 mice each. Bars represent SEM; statistical analysis was performed using two-sided unpaired t test. (F) Quantification of ASC by ELISPOT, plates were coated with WT-HA and MUT-HA, and ASC were quantified in mln of WT vs MUT infected mice. Spot-forming cells were normalized to 10^6^ cells. Data from 5 mice per group. Bars represent SEM, statistical analysis was performed using two-sided unpaired t test. (G) Serum reactivity to WT-HA and MUT-HA in WT- and MUT-infected mice at 14 days post infection. Area under the curve (AUC) quantification is depicted. Data represent four independent experiments with 5 mice each (n = 40). Bars represent mean ± SEM; statistical analysis was performed using a two-sided unpaired t test.

Remarkably, ∼50% of the HA-directed GC and almost 100% of the MBC response in WT infected mice was specific for the E10 epitope in mln (Fig. 6E), lungs and spleen (Fig. S5B-S5E). This is an extreme case of immunodominance where the majority of the B cell response was focused on a very small antigenic surface of H7 HA. Importantly, we did not detect any gross differences in GC size between WT and MUT after infection with the two viruses (Fig. S5F). However, in mice infected with MUT virus, this extreme immune focusing was lost, with only approximately 10% of B cells recognizing the mutated E10 epitope (Fig. 6E). The results here suggest a very strong immune focusing on the E10 epitope upon WT virus infection, which was however lost after nanobody-driven mutations.

To verify whether the immunodominance in WT infected mice extended to the antibody secreting cells (ASC) and the Ab compartments, we performed ELISPOT and ELISA. Consistent with B cell results, ASC immunodominance was very pronounced in WT-infected but not in MUT-infected mice (Fig 6F), a result that was further confirmed when analyzing serum Abs (Fig 6G and Fig S5G). Here, more than half of the Ab reactivity was lost when testing WT-infected sera on the MUT HA protein, confirming the strong immunodominance of the E10 epitope. In our study, we have not investigated the immunodominance to this epitope in different animal species, indeed existing data on species specificity of antibody immunodominance is scarce and contradictory<u>^66,69^</u>. However, it is plausible to think that this site may be under selection pressure in its natural host, and despite that, it continues to show remarkable conservation, possibly because of the lower fitness of MUT virus (Fig 5H).

### H7-HA_166-186_ peptide immunization confers partial *in vivo* protection

Having verified the ability of E10 to provide heterosubtypic protection and the relative stability of its epitope, we decided to verify whether immunization with peptide containing the E10-binding motif would be sufficient to provide protection in animals. Therefore, we expressed the 21 mer peptide corresponding to H7HA_166-186_ sequence (KSYKNTRKSPAIIVWGIHHSV) linked with OVA in the C-terminal to increase immunogenicity and availability of T cell epitopes. The peptide corresponds to two β-sheet in HA1 (Fig 7A). We immunized mice thrice, collected serum 6 days after the third immunization and challenged the mice with a lethal dose of H7 14 days after the last vaccination (Fig 7B). Serum Abs from mice vaccinated with the peptide showed a remarkable breadth, binding not only full length H7 HA but also H3 and H1 HAs, in line with E10 broad reactivity (Fig 7C and Fig S6). The binding capacity was confirmed also by WB analysis of viral lysate (Fig 7D). To check the ability of serum to recognize live virus, we tested sera from peptide-immunized mice for their ability to block hemagglutination: similar to E10, peptide-immunization elicited Abs capable of potently block hemagglutination of viruses belonging to both groups (Fig 7E). Importantly, peptide immunization slightly delayed weight loss and protected 60% of the vaccinated mice against H7 viral challenge, confirming the central role of this epitope in protection (Fig 7F).

**Figure 7.**
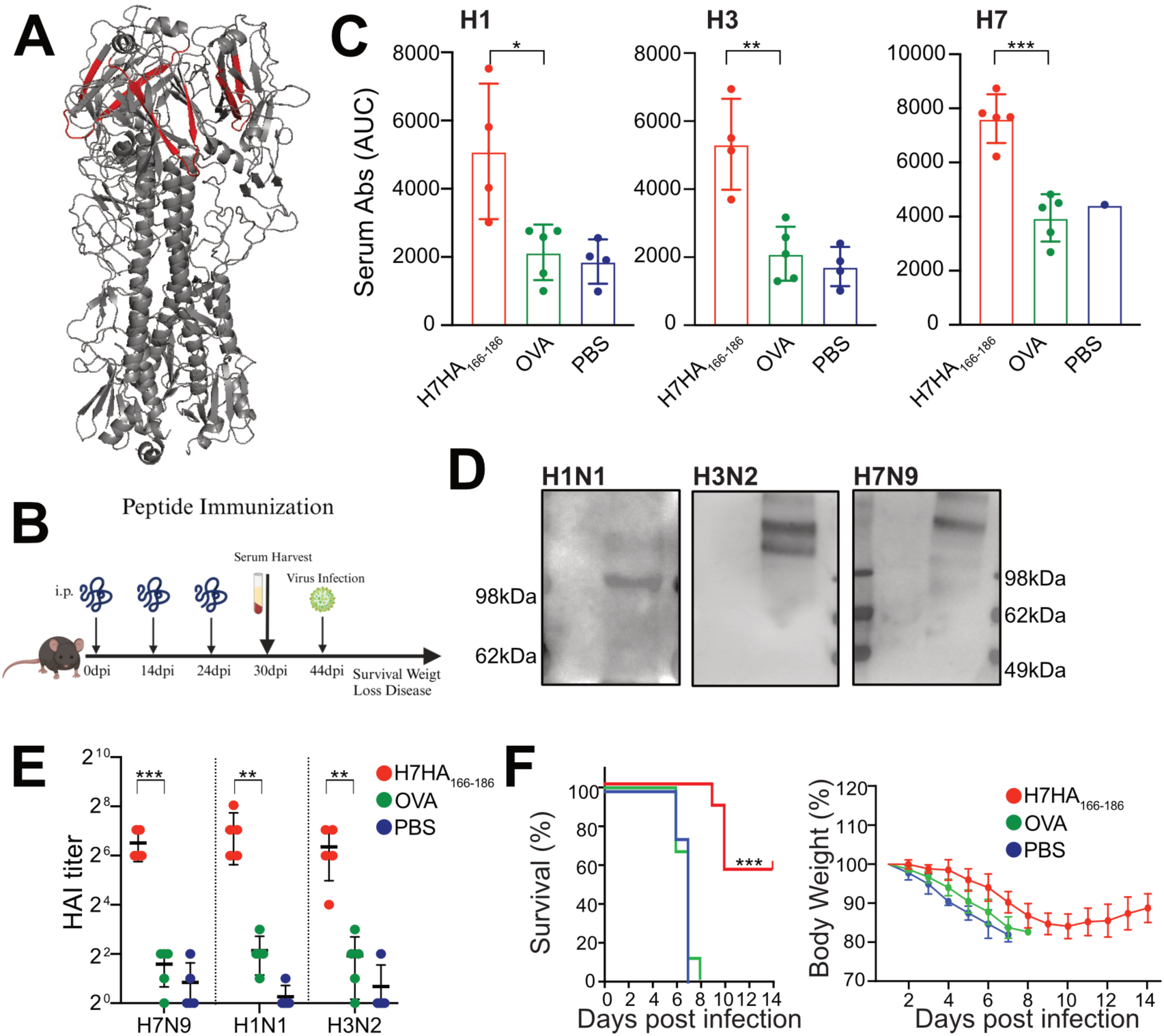
H7HA_166-186_ peptide immunization confers partial protection. (A)Schematic illustration of H7HA166-186 peptide (shown in red) within the HA protein timer (gray) and characteristic of H7HA_166-186_ peptide. (B) Schematic timeline of peptide immunization protocol. (C) Serum binding analysis of H1N1/H3N2/ H7N9 HA protein using serum from mice immunized with H7HA_166-186_ peptide, OVA peptide and PBS. Area under the curve (AUC) quantification of the curves depicted. Data represent two independent experiments with 4-5 mice per group. Bars represent mean ± SEM, statistical analysis was performed using a two-sided unpaired t test. (D) WB analysis of HA protein recognition from MDCK cells infected with H1N1, H3N2, or H7N9 viruses, using serum from H7HA166-186 peptide-immunized mice. First antibody: serum from H7HA_166-186_ peptide-immunized mice, second antibody: anti-mice IgG. (E) Hemagglutination inhibition assay (HAI) tests the binding ability of H7HA_166-186_ immunized serum with H1N1/H3N2/ H7N9 viruses. Data represent the means of two independent experiments. (F) Weight loss and survival curves of mice immunized with H7HA_166-186_ peptide. Mice (n = 10 per group, two independent experiments) were infected with H7N1 (1 TCID_50_, lethal dose) after being immunized three times with H7HA_166-186_ peptide at day 42. As shown in the schedule in 7B.

Overall, peptide immunization, despite its notorious limitations^70^, was able to elicit significant levels of cross-reactive Abs in mice and to afford partial protection against lethal viral challenge thus highlighting the potential and relevance of E10 epitope as vaccine target.

## Discussion

H7N9 remains a serious public health threat with frequent human infections, which are often lethal<u>^12^</u>. Furthermore, the risk of virus adaptation, gaining human-to-human transmission ability, in a naïve population, is a looming danger. Developing novel therapeutic tools to treat zoonotic infections but also discovering sites of vulnerability on this virus is imperative. Compared to mAbs, nanobodies represent a promising new avenue for preventing and treating influenza due to their superior characteristics and distinct binding capacity compared to conventional antibodies<u>^40^</u>. Here, we isolated a broadly neutralizing nanobody, E10, capable of binding multiple strains of influenza A virus. To enhance the therapeutic potential of E10, we added human Fc fragment, improving its structural similarity with human antibodies and affording it with the ability to trigger effector functions. The neutralizing mechanism of E10 involves blocking viral attachment of the HA protein. E10 had remarkable *in vitro* and *in vivo* breadth, providing neutralization and conferring protection against group 2 (H7, H3) and group 1 (H1) strains.

We utilized multiple methods to identify E10 binding site and pinpointed the area around residues K166 and S167 as its binding site. This area included two conserved-beta sheets and was part of the “lateral patch” of HA. Because of antigenic drift, the HA head domain is highly variable among different strains<u>^71^</u>. However, a few regions in the head have gained traction as potential targets of broadly neutralizing Abs. One is the receptor binding site (RBS)<u>^72^</u>, with Abs targeting this area mostly interacting with highly conserved residues of the RBS by inserting their HCDR3<u>^60^</u>. In addition, lateral patch” binding antibodies have been receiving more and more attention<u>^5,27,29,73-75^</u>. For instance, CL6649 can recognize the lateral patch and binds most of the H1N1 viruses ranging from 1977 (seasonal) to 2012 (pdm2009)<u>^29^</u>. Furthermore, Fab6649 also recognized the same area in H1 viruses<u>^29^</u>. Finally, a recent study reported identification of two human H7N9 mAbs targeting the lateral patch on HA head<u>^5^</u>. These could bind to several H7 strains but also H10 and H12, which are close phylogenetically. Importantly, none of these Abs are able to cross-react between different HA groups or even distant types, making E10 a remarkable nanobody. Indeed, cross-group recognition and neutralization is a rare but desirable feature of anti-HA Abs. Possibly, thanks to the unique feature of alpaca nanobodies we increased the likelihood of targeting conserved epitopes in a unique way.

Moreover, lateral patch polyclonal Abs are also induced by pandemic H1 vaccination, though it remains unclear whether these antibodies can cross-react with other influenza subtypes<u>^73^</u>. From the same study, the authors isolated human mAbs and identified a crucial, conserved Y-x-R or Y-R motif within the H-CDR3 region<u>^73^</u>. In addition, these mAbs mostly showed VH3-23/Vk1-33 usage. Among those KPF1, contains a Y-x-R motif within the H-CDR3<u>^22,73^</u>. Notably, our E10 nanobody possesses the “YCSFR” sequence in its CDR3, which is absent in other nanobodies we have identified.

In our study we also demonstrate that peptide immunization is also able to generate cross-reactive Abs that can neutralize *in vitro* and partially protect *in vivo*. The partial *in vivo* protection should not discourage as peptide immunization is notoriously poor at eliciting Abs<u>^70^</u>, it however provides an excellent blueprint on the immunogenicity and potency of this site which could be exploited in future studies with more advanced computational immunogen designs<u>^76^</u>.

With H5N1 infections on the rise, research on animal viruses is an urgent priority and H7N9 presents a significant pandemic risk. Here, besides the identification of the nanobody we also tested basic immunodominance characteristics of the virus. Immunodominance for H7 has not been extensively studied and defining it is not a mere academic exercise but may inform on virus adaptation and mutation potential. Indeed, immunodominance has been suggested as potential driver of antigenic drift<u>^20^</u>. Several studies have now pointed out that certain numbers of individuals have a very focused Ab response, and these may be driving antigenic drift<u>^20,64-67^</u>. Generally, in mice, immune response to IAV is broad and H1N1 virus require over 10 mutations to fully escape the serum of infected animals<u>^50^</u>. Remarkably we show that mutations in the E10 epitope almost completely abolish HA recognition by GC and MBC in mice. Despite this, the residues remain conserved among most of the circulating H7 strains encouraging potential therapeutic use. This suggests either a different immunodominance pattern with low immune pressure in avian hosts or poor fitness of the escape mutants. Whether immunodominance in different hosts is conserved is debated<u>^66,69^</u>, but some data points towards a conserved hierarchy of the response between avian and mammal hosts<u>^69^</u>. Similarly, the ease of isolation of mAbs targeting a similar site in humans^5^ suggests this epitope to be under constant immune pressure. However, our *in vivo* and *in vitro* data suggest this epitope to be needed to maintain a strong viral fitness and therefore, while escape mutants may emerge, they are unlikely to be able to compete with WT virus. Obviously increased immune pressure may favor escape in this site, as demonstrated for other conserved Ab epitopes<u>^77-79^</u>, however, in addition to showing lower viral fitness, the MUT virus showed also increased epitope spread after infection, favoring a broader immune response. This is yet another good news for the potential use of E10 as therapeutic nanobody or for the targeting of the epitope with vaccine constructs.

Despite the limitations in our experimental setup, such as the lack of definitive structural determination by cryo-EM and Biolayer Interferometry (BLI), which could have provided more detailed insights into the antibody-HA interaction and affinity, we successfully mapped the binding area of E10 as a conserved β-sheet on the HA surface. Overall, E10 may add to our arsenal of treatments for zoonotic infections and its epitope and associated peptide may provide a blueprint for future universal influenza vaccine development.

## Material and method

### Inclusion and ethical statement

All the studies were constructed according to ethical permits 1666/19, 38/23 granted by the Gothenburg Regional Animal ethics committee, as well as the Guide for the Care and Use of Laboratory Animals of the Ministry of Science and Technology of the People’s Republic of China. Experiments involving H7N9 avian influenza viruses were performed in a biosafety level 3 laboratory, approved by the Chinese Ministry of Agriculture and Rural Affairs. The protocol was approved by the Committee on the Ethics of Animal Experiments of the Harbin Veterinary Research Institute (HVRI) of the Chinese Academy of Agricultural Sciences (CAAS). Details regarding the facility and biosafety measures have been previously reported<u>^80^</u>.

### Animals

Mice, Female C57BL/6 mice, aged 8-12 weeks, were purchased from Janvier, France, and housed in a specific pathogen-free (SPF) facility at f the Experimental Biomedicine Unit, University of Gothenburg. Additional female C57BL/6 mice were obtained from SPF (Beijing) Biotechnology Co., Ltd. in China and housed under similar conditions. Alpaca, male, aged 2.5 years, were provided by a local farm in Gansu Province, China.

### Cells, viruses and plasmids

HEK293T (American Type Culture Collection, ATCC, CRL-3216), MDCK (ATCC, CCL-34), MDCK-SIAT1 (created and gifted by Dr. Jonathan Yewdell at NIH) cells were cultured in DMEM (Gibco, C11995500BT) with 10% (vol: vol) FBS (Gibco, 10,270–106) and penicillin-streptomycin (Gibco, 15,140,163). A549 cells (ATCC, CCL-185) were maintained in Kaighn’s modified Ham’s-F12 medium (Gibco, C11330500BT) with 10% FBS and 0.01% penicillin-streptomycin. All these cells were cultured and maintained at 37°C with 5% CO_2_. HEK293F (ATCC) was grown in Freestyle 293 Expression Medium (Gibco, 12338018) with 0.01% penicillin-streptomycin and cultured at 37°C with 8% CO_2_ at 125 rpm.

A/Environment/Suzhou/SZ19/2014(H7N9)(SZ19) was isolated and stored in our laboratory, A/Puerto Rico/8/1934 (PR8, H1N1) was stored in our laboratory<u>^53^</u>. A/Puerto Rico/8/1934 (PR8, H1N1), influenza A virus A/Hongkong//1968 (X31, H3N2) was stored in our laboratory in Gothenburg. H7N1 virus, H7N1-MUT virus (HA comes from SZ19, others come from PR8) were rescued in our laboratory in Gothenburg.

The H7-HA and H7-HA-K166T, S167L were constructed containing the H7 ORFs: one for the H7N1_K166, S167_ and another for the H7N1_K166T, S167L_ mutated influenza A/Environment/Suzhou/SZ19/2014/H7N9 strains, labeled as WT and MUT, respectively. The cloning strategy involved double-digestion of the pDZ plasmid with E*co*RI-HF (NEB, R3101S) and X*ho*I (NEB, R0146S), followed by In-Fusion cloning (Takara Bio, 638948) with three PCR-amplified fragments, containing the H7 ORF and its flanking regulatory regions. Primer sequences are shown in S1 Table.

### Sequencing

Constructed plasmids were sequenced and analyzed by Eurofins and Tsingke Biotechnology Co., Ltd.

### Alpaca immunization and phage display selection

Immunization: Two years and half male alpacas were immunized five times, with 14 days interval, with inactivated (0.06% formaldehyde) and purified H7N9 virus (50 µg/dose). The first immunization used Freund’s Complete Adjuvant (FCA, Thermo, 77140) followed by four subsequent immunizations with Freund’s Incomplete Adjuvant (Thermo, 77145). The indirect ELISA method was used to measure the IgG titer against H7N9, initiating phage display construction when the titer exceeded 1:64000.

Phage Display: After 14 days from the last immunization, peripheral blood mononuclear cells (PBMC) were isolated from whole blood of alpaca, and RNA was extracted to synthesize cDNA. VHH fragments were amplified via nested PCR using specific primers, then cloned into the pComb phage vector and transformed into E. coil SS320 competent cells. The positive clones were sequenced, and then analyzed the diversity of the antibody library with the calculated antibody library capacity, the remaining culture was transferred to Amp-Kan medium for expansion culture, and PEG/NaCl was used to concentrate to obtain the original antibody library. ELISA plates were coated with inactivated purified H7N9 at 1 μg/well, 1% polyvinyl alcohol (1% PVA) was set as a no-antigen control, 100 μL of the original antibody library was added to each well, incubated at room temperature for 2 h, and the solution was discarded; 0.1 mol·L-1 HCl was added for elution for 5 min, and an equal volume of Tris-HCl was used for neutralization; the eluate was added to logarithmic phase NEB5aF’ bacteria, cultured at 200 r/min, 37 ℃ for 1 h, and M13 assisted phage rescue for 1 h. An appropriate amount of bacterial solution was taken for 10-fold dilution and titrated on the plate; the remaining bacterial solution was cultured overnight. Enrichment and panning were respect three times and selected by ELISA with anti-mouse M13 antibody. Then select the positive bacteria and express the nanobodies to pcDNA13.1-Fc vector and transform to DH5α competent cells. After three rounds, 96 colonies were selected, and the eluted phages were further characterized for binding by indirect-ELISA. Primer sequences are shown in S1 Table.

### Nanobody production and characterization

For protein production, nanobodies were expressed in HEK293F cells maintained in Expi Expression Medium (Gibco, 12338018), followed by transfection using the ExpiFectamine™ 293 Transfection Kit (Gibco, A14525). After 5 days, culture supernatants were filtered and purified using Protein G columns on an FPLC AKTA start system.

### Confocal microscopy

HEK293 and A549 cells (5◊10^5^ cells/well) were plated on coverslips. Cells were allowed to attach for 8h and were left uninfected or were infected with H1N1/H3N2/H7N9 IAV (MOI=0.1) for 24-36h in serum free media. Cells were fixed with 4% paraformaldehyde for 20 min at room temperature and washed with PBS three times. Cells were permeabilized with 0.1% Triton X-100 in PBS for 10min, then blocked with 5% skimmed milk for 1h. Then cells were incubated with nanobodies as the primary and anti-human IgG (Fc specific)-488 (Invitrogen, A55747) as secondary antibodies and DAPI (Beyotime, C1002). Leica microscope (TCS SP8) were used to observe the strained cells with a 100◊ oil objective.

### Western blotting

Protein samples (5-20 µg) were separated into 4-12% Bis-Tris gels, transferred to Nitrocellulose membrane (Cytiva, 1060000). Proteins of interest were analyzed by hybridization with their corresponding antibodies (anti-GAPDH (ab181602), anti-NP (Sino Biological, 11675-V08B)), horse anti-mouse IgG (Vector laboratories, ZK0403), Mouse Anti-Human IgG Fc Antibody (GenScript, 50B4A9) and visualized by chemiluminescence using Thermo Scientific SuperSignal™ West Dura Extended Duration Substrate (ThermoFisher, 34076).

### Neutralization assay

Serial dilutions of nanobodies (start from 100ng/µL) or mouse serum were mixed with 100 TCID_50_ of virus (H7N9/H3N2/H1N1) for 1 h at 37°C. The mixture was then added to cultured MDCK cells in 96-well plates for 1h. After PBS washed 3 times we incubated with DMEM containing 1 μg/ml TPCK-treated trypsin and 0.01% penicillin-streptomycin at 37 °C. After 20h viral infection cells was quantified by indirect ELISA with nanobodies against the nucleoprotein (NP) of influenza A virus (Sino Biological, 11675-V08B). The final concentration of nanobody that reduced infection to 50% (IC_50_) was determined using GraphPad Prism 6 software. Or after 72h incubation we took supernatant and added with 1% chicken red blood cells (RBC), then calculated the 50% neutralization titer. Log50 % neutralizing titer = *L*-*d* (*s*-0.5), where *L* is the log of the lowest dilution factor, *d* is the difference between the dilution factors, and *s* is the sum of the ratios of positive wells.

### Recombinant hemagglutinin production and testing

Recombinant HA from A/Environment/Suzhou/SZ19/2014(H7N9) (SZ19), A/Puerto Rico/8/1934 (PR8, H1N1), Hongkong//1968 (X31, H3N2) and SZ19-HA-K166T, S167L were expressed and purified as previously described. Briefly, proteins were expressed in 293F cells and purified by affinity chromatography followed by size exclusion.

### Enzyme-linked immunosorbent assay (ELISA)

For serum binding test, 96 well plates were coated with recombinant protein (H7-HA (WT-HA) or H7-HA-K166T, S167L (MUT-HA)) at 1 μg/ml in PBS and incubated overnight at 4°C. Wells were blocked with 100 μL 2% BSA in PBS for 1h at RT. Plates were washed three times with PBS + 0.05% Tween and incubated with two-fold serially diluted sera (starting from 1:100) in PBS with 0.05% Tween for 1.5h at RT. After three washes, anti-mouse Igκ (Sino Biological, 68077-R008-H) at a 1:1000 dilution was added, and plates were incubated for 1h at RT. After three washes, plates were developed using TMB (Thermo Fisher, catalog: 34029) for 5 minutes at RT and subsequently blocked with H_2_SO_4_. The absorbance was measured with a TECAN Sunrise absorbance microplate reader (catalog: 16039400) at 450 nm.

For nanobody binding test, we coated 96-well plates with different strain of UV-inactivated virus (H1H1/H3N2/H7N1) or different strain of HA recombinant protein (H1N1-HA/H3N2-HA/H7N9-HA) at 1 μg/ml in PBS and incubated overnight at 4°C. Wells were blocked with 100 μL 2% BSA in PBS for 1h at room temperature. Plates were washed three times with PBS + 0.05% Tween and plates were incubated with two-fold serially diluted nanobodies (starting from 100ng/mL) in PBS + 0.05% Tween for 1.5h at RT. After three-time washes, mouse anti-human IgG Fc Antibody (Sino Biological, SSA001) at 1:6000 dilution was added, and the same as above.

### Hemagglutination assay

Serial dilutions of nanobodies (start from 100ng/µL) were mixed in equal volumes with 100 TCID_50_ of virus (H7N9/H3N2/H1N1) for 1 h at 37°C. The mixture was then added to cultured MDCK cells in 96-well plates for 1h. After PBS washed 3 times, we incubated with DMEM containing 1 μg/ml TPCK-treated trypsin and 0.01% penicillin-streptomycin at 37 °C for 72h. Viral infection was quantified by hemagglutination assay. Briefly, 50 µL virus supernatant were incubated with 1% chicken RBC suspension in a round-bottomed 96-well plate. And incubated at room temperature for 1h before the results were inspected.

For virus rescue detection, the hemagglutination assay was performed using 1% chicken RBC suspension to confirm virus rescue and neutralization titer. Briefly, 50 µL virus supernatant was used to make 2× serial dilutions in a round-bottomed 96-well plate. The diluted virus supernatants were mixed with 25 µL of 1% chicken RBC in each well and incubated at room temperature for 1h before the results were inspected.

### Detect nanobody mechanism to entry into cells

For nanobody attachment detection, H7N9 virus (MOI=10) was incubate at 37 ℃ for 1h. Then added into A549 cells and kept in 4 ℃ for 1h, andput in 37 ℃1h, then collected cells for RNA extraction. For nanobody internalization detection, H7N9 virus (MOI=10) was incubate at 37 ℃ for 1h. Then added into A549 cells and kept in 4 ℃ for 1h, then washed with pH=3 PBS three times and put in 37 ℃ for 4h incubation. Then cells were collected for RNA extraction. For nanobody replication detection, H7N9 virus (MOI=10) was incubate at 37 ℃ for 1h. Then added into A549 cells and kept in 4 ℃ for 1h, then washed with pH=3 PBS three times and put in 37 ℃ for 8h incubation. Then cells were collected for RNA extraction. For nanobody release testing, H7N9 virus (MOI=10) was incubate at 37 ℃ for 1h. Then washed with pH=3 PBS three times. After 10h washed with PBS three times and added nanobodies for 2h and collected cells for RNA extraction.

### RNA isolation and quantitative RT-PCR

Total RNA from cells was extracted with TRIzol following the manufacturer’s instructions. For mRNAs, total RNA was transcribed into cDNA using M-MLV Reverse Transcriptase, according to the manufacturer’s protocol (Promega, M1701). GAPDH was used as an invariant control for mRNAs. Real-time PCR was carried out using the Light Cycler 480 System (Roche). The RNA level of each gene was shown as the fold of induction (2^−ΔΔCT^) in the graphs. Primer sequences are shown in Table S1.

### Prophylactic and therapeutic efficacy in mice

For prophylactic evaluation, groups of 16 mice were intraperitoneally (i.p.) injected with 10 mg/kg purified nanobodies 24h before being challenged intranasal (i.n.) with 25 µL of virus (H7N9:10^6^ EID_50_, H3N2:10^7^ TCID_50_, H1N1: 3×10^3^ TCID_50_) suspended in Hanks’ Balanced Salt Solution (HBSS) + 0.1% fetal bovine serum (FBS). Mice were anesthesia with carbon dioxide ice or isoflurane during virus challenge. For therapeutic assessment, mice were infected i.n. with virus (H7N9: 10^6^ EID_50_, H3N2:10^7^ TCID_50_, H1N1: 3×10^3^ TCID_50_, H7N9-MUT (T166, S167): 10^6^ EID_50_ and treated 2h later with a single i.p. dose of 25 mg/kg purified nanobodies. Control groups received PBS instead of nanobodies. Mice were monitored daily for clinical signs and body weight loss. On predetermined days post-infection, mice were euthanized, and tissues (nasal turbinate, lung, spleen, kidney, brain, liver) were collected for viral titration and histopathological examination.

### Hemagglutination inhibition (HAI) assay

H7N9 viral stocks were standardized to 4 HA units using 1% chicken RBCs before use in HAI assays. Serial two-fold dilutions of 25µL nanobodies (starting at 300ng/µL) were prepared in V-bottom microtiter plates. Subsequently, 25µL of the 4 HA unit virus was added to each well and incubated for 30 min at room temperature (RT). After incubation, 50μl of 0.5% chicken RBCs was added, and the mixture was further incubated for 30 min at RT. HA inhibition was visually assessed by the formation of well-defined RBC “buttons” or teardrop patterns upon plate tilting. HAI titers were reported as the reciprocal of the highest dilution that completely inhibited hemagglutination.

### Escape mutation selection

For egg-based selection,100µL of H7N9 virus (10^6^ TCID_50_) was incubate at 37℃ for 1h with 100µL of nanobodies at varying concentrations (500, 50, 5, 0µg/mL). The virus-antibody mixture was then injected into 10-day-old SPF (specific-pathogen-free) eggs, and after 40-44h incubation, the supernatant was collected and tested by hemagglutination assay (HA). Then detected the existence of escape mutation by HA and HAI assays. Virus samples positive in both assays were sequenced using next-generation sequencing analysis (NGS) to identify amino acid mutations in the HA segment.

For cell-based selection, MDCK cells were infected with H7N9 virus in the presence of increasing concentration of nanobodies. After 1h at 37°C with 5% CO_2_, the virus containing medium was replaced with virus growth medium (VGM) containing nanobodies, and incubation continued until cytopathic effect (CPE) was observed. Supernatants were collected, and the virus was passaged 10 times, doubling the nanobody concentration after each passage. Viruses from the final passage were sequenced using NGS to identify mutations across all segments.

### Next-generation sequencing (NGS) analysis

Viral RNA was extracted from the egg supernatant and reverse-transcribed into cDNA using the Uni12 primer (5’-AGCRAAAGCAGG-3’). Full-length HA gene amplification was conducted using specific primers (Supplementary Table S1). NGS was performed and analyzed by Tsingke (Tsingke Biotechnology Co., Ltd.).

### Phage display mapping of mAbs

To map the nanobody epitopes, a phage display library was constructed. The full-length H7 protein was divided into 20-mer sequences overlapping by 5 amino acid. The individual plasmids were transformed in ElectroMAX DH12S Cells (Invitrogen, catalog: 183812017) to create the library. The library was inserted into phages by using an additional helper phage and the number of phagemid particles was determined by infecting *E. coli* and counting colonies.

For epitope binding analysis, 96-well plates were coated with each nanobody and blocked with BSA. Phagemid-carrying particles were added and after incubated, unbound particles were washed off, while bound phage were eluted, neutralized and stored. Phage suspensions were used to identify binding epitopes, and individual colonies were sequenced to map the epitopes on a 3D structure using PyMOL 2.5.5. The entire library was also sent for Illumina sequencing to Eurofins Scientific to make sure that each peptide was present in the library.

### Viral growth kinetics

Triplicate wells of confluent MDCK cells were infected with WT and MUT virus at a MOI= 0.001 and incubated with BSA-MEM containing 1 μg/ml TPCK-treated trypsin at 37 °C. Supernatants were harvested at 6-, 12-, 24-, 36 h post-infection. Virus titers were determined using 1% chicken Red Blood Cells with the supernatant of MDCK cells in 96-well plates after incubated with different harvested supernatant dilution 48h.

### Rescue of H7N1 virus

The H7N1-WT and H7N1-MUT viruses were rescued under BSL-2 conditions, as viruses containing HA of Influenza A/Environment/Suzhou/SZ19/2014 and the backbone segments of A/Puerto Rico/8/1934/H1N1 (PR8 strain).

Influenza A reverse genetics was performed using a previously established protocol<u>^81^</u> to rescue the H7N1-WT and H7N1-MUT viruses. Briefly, plasmid cocktails were prepared to contain pDZ plasmids of the seven PR8 backbone segments with either the pDZ H7 SZ19-WT or the pDZ H7 SZ19-MUT plasmid respectively. Co-cultures of HEK293T and MDCK-SIAT1 cells were reverse transfected with the respective plasmid cocktails using TransitLT1 reagent (Mirus Bio, lyec-1) in a 1:4 ratio. Monolayers of MDCK-SIAT1 cells were then infected with the co-culture supernatant and observed for cytopathic effect after 72h. The virus rescues were confirmed using hemagglutination assay and their H7 ORF sequences were confirmed using Sanger sequencing.

### HA sequence analysis and alignment

To determine the distributions of amino acids at position 166,167 of HA in influenza A virus derived from different species, a total of HA amino acid sequences was downloaded from the Global Initiative on Sharing All Influenza Data (GISAID) (https://www.gisaid.org/) database and the Influenza Virus Database of GenBank (https://www.ncbi.nlm.nih.gov/genomes/FLU/Database/nph-select.cgi?go_database). The amino acid sequences of each HA were aligned using MAFFT<u>^82^</u>. Base compositional data of the amino acid at position 166,167 were then graphically plotted using the Python language.

### Virus infection of mice and flow cytometry analysis

The rescued virus H7N1 (WT) and H7N1-T166, L167 (MUT) were generated following previous protocols. C57BL/6 mice, maintained in our laboratory in Gothenburg, were infected intranasally with either WT virus (0.1 TCID_50_) or MUT virus (50 TCID_50_) per mouse in Hanks’ Balanced Salt Solution (HBSS) + 0.1% fetal calf serum (BSA) intranasally in a volume of 25μL/mouse. 14 days post-infection, the mice were sacrificed and harvestmediastinal lymph nodes (medLNs), spleens and lungs for further analysis. The harvested organs were proceeded, and red blood cells were lysed using ACK lysing buffer to obtain a single-cell suspension. For B cell characterization, 25μL of pre-mixed extracellular antibody mixture was added to each sample and incubated for 30 minutes at 4°C. The following antibodies were used for extracellular B cell staining: 1:700 anti-mouse CD3 BV510 (BD, catalog: 563024), 1:200 anti-mouse IgM PE/Dazzle 594 (BioLegend, catalog: 314529), 1:200 anti-mouse IgD BV785 (BD, catalog: 563618), 1:200 anti-mouse CD38 FITC (BioLegend, catalog: 102705), 1:200 anti-mouse GL7 PE (BioLegend, catalog: 144607), and 1:200 anti-mouse B220 Pe-Cy7 (BioLegend, catalog: 103221). Following the antibody incubation, the samples were washed with 1mL of FACS-buffer (DPBS + 2% FBS + 0.5mM EDTA), and the supernatant was discarded. To assess HA-specific B cells, 100μL of premixed WT/MUT HA proteins were added to each tube and incubated for 1h at 4°C. WT-HA protein was conjugated with streptavidin-APC (Bioligand, catalog: 405243), Brilliant Violet 421 Streptavidin (BioLegend, catalog:405225), MUT-HA protein was conjugated with Brilliant Violet 650 Streptavidin (BD, catalog:563855), Percp5.5 Streptavidin (BioLegend, catalog:405214) before the experiment. The samples were then washed and incubated for 30 minutes at 4°C. To exclude dead cells, Live/Dead Aqua (Invitrogen, catalog: L34966) staining was performed, followed by fixation of the cells with 1.5% paraformaldehyde (PFA). After fixation, the samples were washed, resuspended in 200μl of FACS buffer, and stored at 4°C until analysis.

### B cell ELISpot assay

Mice were sacrificed 14 days after being infected with WT (0.1 TCID_50_) /MUT (50 TCID_50_) virus 14 days, and their mediastinal lymph nodes (medLNs) were harvested and processed for analysis. 96-well ELISpot plates (Millipore Sigma) were coated with 100 μL per well of recombinant H7-HA protein and H7-HA-T166, L167 at a concentration of 1 μg/mL, and incubated overnight at 4 °C. The next day, the plates were blocked with 100μL of PBS containing 2% BSA and incubated for at least 1h at RT. Cells isolated from medLNs were counted using a Muse cell analyzer (Merck Millipore) in 0.1% BSA in PBS. Each well was seeded with 1.5×10^5^ cell in 150μL of DMEM supplemented with 10% FBS and 10μg/mL gentamicin, then serially diluted 3-fold and incubated overnight at 37°C. Plates were washed three times with PBS containing 0.05% Tween, followed by a 2h incubation with 50μl anti-mouse IgG H+L HRP (Aviva Systems Biology, catalog: ORA 04973) at RT. After three additional washes, the plates were developed using BD ELISpot AEC Substrate solution and incubated in a humid chamber for 10 minutes at RT before stopping the reaction. A CTL ImmunoSpot plate reader was used for imaging, and spots were manually counted. The number of spot-forming cells was normalized to 10^6^ cells.

### Atomic model building and refinement

For structure determination, a model of nanobodies was generated using ImmuneBuilder<u>^54^</u>. The nanobodies link with Fc tag was AlphaFold 3.0<u>^83,84^</u>, SZ19-HA and SZ19-HA-_T166,L167_ were generated by Swiss-model (https://swissmodel.expasy.org/). Structures were analyzed and figures were generated using PyMOL (http://www.pymol.org). Final model statistics were summarized in supplementary Fig. S1B.

### Peptide preparation and immunization

H7-HA_166-186_ peptide (length: 21 amino acids) were linked with OVA tag at the C-Terminal modification and conjugation on C terminal-COOH. The sequence of H7-HA_166-186_ peptide is “KSYKNTRKSPAIIVWGIHHSV”. Mice were immunized with H7-HA_166-186_ /OVA peptide or just PBS three times with i.p. The first two times each mouse immunized with 100µg H7-HA_166-186_ /OVA peptides in 100µL PBS and mixed with 100µL equal value of AddaS03™ Adjuvant (InvivoGen, vac-as03-10). The third time peptides were same with the first two times just without adjuvant. The H7-HA_166-186_ peptide was synthesis by GenScript and the purify is ≥95%.

### Statistical analysis

GraphPad Prism software was used for statistical analysis. To compare the counts of cells between the groups in the ELISpot screening analysis, a one-way ANOVA was employed, followed by Dunnett’s multiple comparisons test to adjust for comparisons between individual pools and the control peptide. For nanobody detection, a two-sided unpaired Student’s t-test was performed. Statistical significance was indicated as follows: **** for p-value ≤ 0.0001, *** for p-value≤ 0.001, ** for p-value ≤0.01, and * for p-value ≤0.05. Additionally, a two-way ANOVA with post hoc Tukey’s HSD test was performed, using an alpha level of 0.05 to identify significant differences across multiple factors. Statistical significance for these analyses was similarly indicated with **** for p-value of ≤0.0001 and *denotes a p-value of ≤0.05.

## Acknowledgments

We would like to thank all members of the Angeletti lab for helpful discussions regarding the project. We thank Zhengxiang Wang for the help on the quantify of virus EID_50_. We would like to acknowledge Protein Production Sweden for provisioning of facilities and experimental support, and we would like to thank M. Andersson and M. Bäckström for assistance. Protein Production Sweden is funded by the Swedish Research Council as a national research infrastructure. We thank the laboratory for Experimental Biomedicine (EBM) at the University of Gothenburg for assistance with mouse breeding and husbandry. The study was supported by grants from the National Key R&D program (grant no. 2021YFD1800204 to Q.Z.), the National Natural Science Foundation of China (grant no. U23A20243 and 32272972 to Q.Z.), the European Research Council (ERC-StG, B16 DOMINANCE, grant no. 850638 to DA); the Swedish Research Council (grant no. 2021-01164, 17 2021-01165 to DA); the Knut and Alice Wallenberg Foundation (grant no 2021.0033 to DA)

## Author Contributions

Conceptualization / Project administration: QZ, DA.

Data collection / Data curation: ZSC, XW, KS, YJ, DFB.

Methodology / Investigation: ZSC, HCH, KS, ML, YJ, QZ, DA.

Formal analysis: ZSC, HCH, XW, QZ, DA.

Writing - Original draft: ZSC, HCH, AAS, DA.

Writing – Review and Editing: All authors.

Technical Support: AAS, GD.

Supervision: QZ, DA.

Funding acquisition: QZ, DA.

## Data Availability

All data presented in the manuscript is available from the corresponding author upon reasonable request

