## Supplementary Material for "Influenza A Virus H7 nanobody recognizes a conserved immunodominant epitope on hemagglutinin head and confers heterosubtypic protection"

**Contents of this file**

Supplementary Table 1

Supplementary Figures 1-6

**Introduction**

This file provides supplemental information for the main manuscript.

**Supplementary Table 1.** PCR primers used in this study.

| Primer name | Primer sequence |
|---|---|
| **Fragment A-Forward** | ATCATTTTGGCAAAGGAATTCGAGCTCGGTACCCGG |
| **Fragment A-Reverse** | GAGTGTTCATTATGTTTTTGTCACCCTGCTTTTGCTCCCC |
| **Fragment B-Forward** | CAAAAACATAATGAACACTCAAATCCTGGTATTCGC |
| **Fragment B-Reverse** | TTTTTTAGTATTATATACAAATAGTGCACCGCATGTTTCCA |
| **Fragment C-Forward** | TTGTATATAATACTAAAAAACACCCTTGTTTCTACTAATAACCCG |
| **Fragment C-Reverse** | AAAAAGATCTGCTAGCTCGAGCATGCCCGGGTACCAT |
| **IgG-Forward** | GTCCTGGCTGCTCTTCT |
| **IgG-Reverse** | GGTACGTGCTGTTGA |
| **VHH-Forward** | CTACAAATGCCTATGCATCCCAGGTGCAGCTCGTGGAGTC. |
| **VHH-Reverse** | AAACAACTTTCAACAGTGGAGGGGTCTTCGCTGTGGTGCG |
| **pComb-Forward** | AAAGAATATCGCATTTCTTCTTGCATCT |
| **pComb-Reverse** | AGGAGACGGTGACCTGGGTCCCCTG |
| **Nb-Forward** | TCCAGTGTGGTGGAATTCGCCACCAAAAAGAATATCGCATTTCTTC TTGCATCT |
| **Nb-Reverse** | TCTAGACTCGAGTCAGTGATGGTGGTGGTGGTGGTGGTGGTGTGAGG AGACGGTGACCTGGG |
| **GAPDH-Forward** | GTCATTGAGAGCAATGCCAG |
| **GAPDH-Reverse** | GTGTTGCTACCCCCAATGTG |
| **HA-Forward** | ATGAACACTCAAATCCTGGT |
| **HA-Reverse** | TTATATACAAATAGTGCACC |

**A**

| | | | | | | | | | | | | |
|---|---|---|---|---|---|---|---|---|---|---|---|---|
| Pro | 2.5844 | 2.5349 | 2.2234 | 2.4812 | 2.6553 | 2.0399 | 2.1616 | 2.2566 | 2.5813 | 2.6464 | 2.2611 | 2.5904 |
| | 2.3277 | 2.5618 | 2.5059 | 2.1862 | 2.4761 | 2.5564 | 2.2086 | 2.2079 | 2.4225 | 2.4103 | 2.3822 | 2.3129 |
| | 2.2422 | 2.2521 | 2.3624 | 2.4614 | 2.1851 | 2.2816 | 2.5236 | 2.5039 | 2.5006 | 2.4689 | 2.3540 | 2.3173 |
| | 2.4294 | 2.2686 | 2.3537 | 2.4503 | 2.5418 | 2.1935 | 2.3901 | 2.2024 | 2.3786 | 2.3759 | 2.3634 | 2.7421 |
| | 2.3006 | 2.3595 | 2.4405 | 2.4652 | 2.5334 | 2.2572 | 2.4085 | 2.3536 | 2.3847 | 2.3429 | 2.2580 | 2.8712 |
| | 2.4748 | 2.4145 | 2.5453 | 2.5109 | 2.4904 | 2.4416 | 2.4677 | 2.4578 | 2.0823 | 2.5587 | 2.4731 | 2.5530 |
| | 2.4345 | 2.5656 | 2.5248 | 2.5474 | 2.8563 | 2.0292 | 2.5360 | 2.2703 | 2.6137 | 2.5589 | 2.5242 | 2.8171 |
| | 2.6393 | 2.5519 | 2.5447 | 2.5620 | 2.3547 | 2.4623 | 2.6422 | 2.4424 | 2.5308 | 2.6024 | 2.5028 | 2.8671 |
| PVA | 0.2206 | 0.1846 | 0.1691 | 0.2315 | 0.1755 | 0.1986 | 0.1734 | 0.1858 | 0.1787 | 0.1862 | 0.1491 | 0.1939 |
| | 0.1587 | 0.1247 | 0.1596 | 0.1545 | 0.1481 | 0.1659 | 0.1534 | 0.1640 | 0.1473 | 0.1510 | 0.1462 | 0.1721 |
| | 0.1922 | 0.1746 | 0.1620 | 0.2247 | 0.1767 | 0.1518 | 0.1847 | 0.2003 | 0.1627 | 0.1471 | 0.1246 | 0.1466 |
| | 0.1470 | 0.1788 | 0.2007 | 0.1523 | 0.1640 | 0.1845 | 0.2081 | 0.1709 | 0.1778 | 0.1514 | 0.1351 | 0.1590 |
| | 0.1829 | 0.1273 | 0.1594 | 0.1970 | 0.2258 | 0.1459 | 0.1816 | 0.1348 | 0.1599 | 0.1210 | 0.1287 | 0.1576 |
| | 0.1453 | 0.0997 | 0.1389 | 0.1578 | 0.1361 | 0.1691 | 0.1795 | 0.1094 | 0.1413 | 0.1308 | 0.1245 | 0.1554 |
| | 0.1891 | 0.1665 | 0.1428 | 0.1706 | 0.1864 | 0.1295 | 0.1366 | 0.1528 | 0.1337 | 0.1153 | 0.1109 | 0.1533 |
| | 0.1818 | 0.1465 | 0.1438 | 0.1604 | 0.1953 | 0.1320 | 0.1499 | 0.1294 | 0.1719 | 0.1248 | 0.1402 | 0.1478 |
| Pro/PVA | 11.7153 | 13.7319 | 13.1484 | 10.7179 | 15.1299 | 10.2714 | 12.466 | 12.1453 | 14.4449 | 14.2127 | 15.165 | 13.3595 |
| | 14.6673 | 20.5437 | 15.7011 | 14.1502 | 16.7191 | 15.4093 | 14.3977 | 13.4628 | 16.446 | 15.9623 | 16.2941 | 13.4393 |
| | 11.666 | 12.8986 | 14.5827 | 10.9542 | 12.3662 | 15.0303 | 13.6632 | 12.5007 | 15.3694 | 16.7838 | 18.8925 | 15.807 |
| | 16.5265 | 12.6879 | 11.7275 | 16.0886 | 15.4988 | 11.8889 | 11.4853 | 12.8871 | 13.378 | 15.6929 | 17.4937 | 17.2459 |
| | 12.5785 | 18.535 | 15.3105 | 12.5137 | 11.2197 | 15.4709 | 13.2627 | 17.4599 | 14.9137 | 19.3628 | 17.5447 | 18.2183 |
| | 17.0323 | 24.2177 | 18.3247 | 15.9119 | 18.2983 | 14.4388 | 13.7476 | 22.4662 | 14.7367 | 19.5619 | 19.8643 | 16.4286 |
| | 12.8741 | 15.409 | 17.6807 | 14.932 | 15.3235 | 15.6695 | 18.5652 | 14.858 | 19.549 | 22.1934 | 22.761 | 18.3764 |
| | 14.5176 | 17.4191 | 17.6961 | 15.9726 | 12.0568 | 18.6538 | 17.6264 | 18.8748 | 14.7225 | 20.8526 | 17.8516 | 19.3985 |

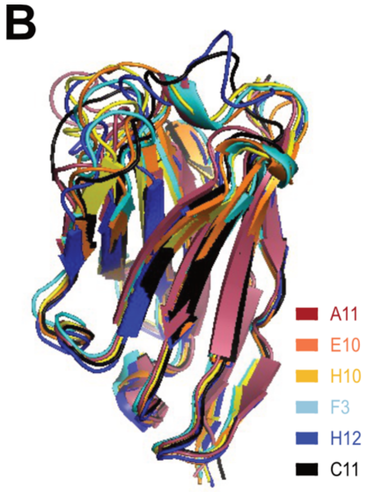


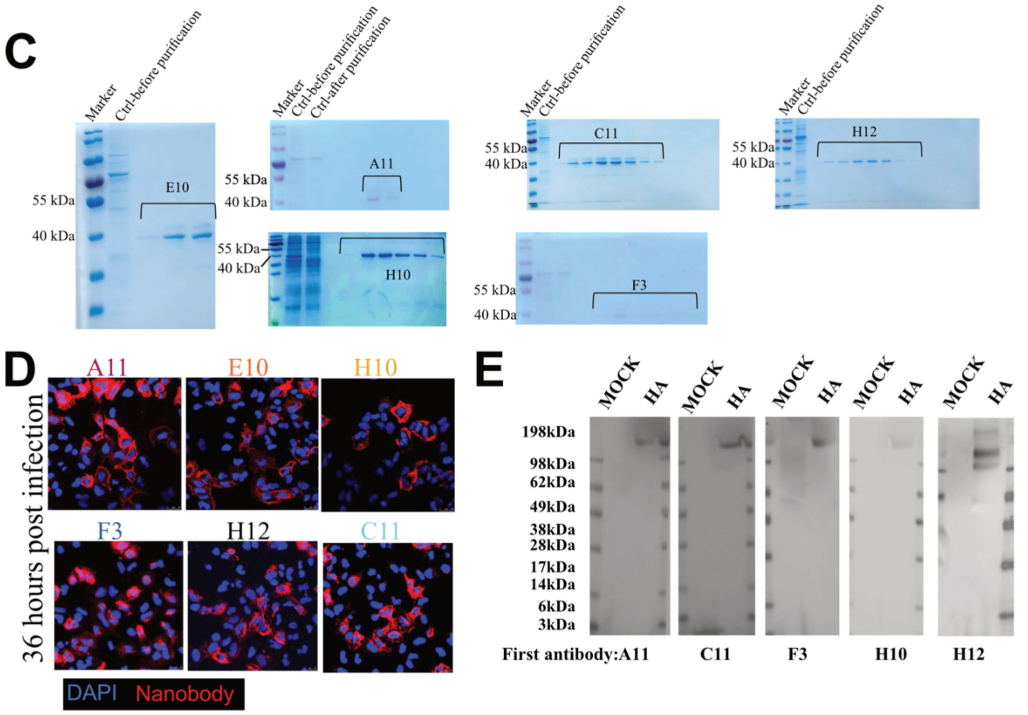


**F**

**Sequence of nanobodies:**

| | FR1 | CDR1 | FR2 | CDR2 |
|---|---|---|---|---|
| **A11** | QVQLVESGGGLVQPGGSLRLSCAAS | GASITTYNMR | WVRQAPGKDLEWVS | QINSDGIRPWYTDSVKG |
| **E10** | QVQLVESGGGLVATGGSLRLSCTAS | GFSLDRYAIG | WFRQAPGKEREGIS | CISSRRDRTNYADSVNGRLT |
| **F3** | QVQLVESGGGLVQPGGSLRLSCAAS | KFTLDNYSIG | WFRQAPGKEREEIS | CSSARDSATYYADSVKG |
| **C11** | QVQLVESGGDLVQPGGSLRLTCALS | LFTTGYYTIG | WFRQAPGKEREGVS | AISYNAGSTDYSDSVKG |
| **H10** | QVQLVESGGGLVQPGGSLRLSCATS | GTTLDNYSIG | WFRQAPGKEHEEIS | CTSAGDGTTYYADSVKG |
| **H12** | QVQLVESGGGLVQPGGSLRLSCAAS | GFTLDYYAMD | YHAIGWFRQAPGKE | REEVSCTSRDGSTNYAESVKG |

| FR3 | CDR3 | FR4 |
|---|---|---|
| RFTISRDNAKNTLYLQMNSLKPEDTAVYYCGN | GAVAGGRLLGG | WGQGTQVTVSS |
| VSRDNAKNTVYLQMNSLKPEDTAVYYCAA | DRSSYTYCSFRLSEYDY | WGQGTQVTVSS |
| RFTISRDNAKSTVYLQLNSLKPEDTAVYYCTA | AGSWYCPATPDY | WGQGTQVTVSS |
| RFTISRDNSKNTVYLQMNSLKPEDTAVYYCAA | KGNWYPSLSRSDYDY | WGQGTQVTVSS |
| RYTISRDNAKNTVYLQMNSLKPEDTAVYYCTA | AGSWYCPATPDY | WGQGTQVTVSS |
| RFTISRDNAKNTVYLQMNSLKSEDTAVYYCTA | AGSWYCPASPDY | WGQGTQVTVSS |

**Figure S1. Generation ad characterization of H7-specific nanobodies.**

(A) Binding ability of 96 colons detected by Indirect-ELISA. (20 high binding colons: yellow). (B) Nanobody structure model created using ImmuneBuilder and AlphaFold 2. (Different colors represent different nanobody: A11: red, C11: black, E10: orange,

F3: blue, H10: yellow, H12: dark bule.) (C) Coomassie Brilliant Blue Staining detection of the expression and size of different nanobodies. (D)Immunofluorescence assay (IFA) showing the recognition of the SZ19 virus by different nanobodies. A549 cells were infected with SZ19 (MOI=0.1) for 36h and stained with distinct primary nanobodies, followed by secondary goat anti-human IgG Fc Alexa Fluor™ 488 (A11: red, E10: orange, H10: yellow, F3: blue, H12: dark blue, C11: black.). (E) Western blot analysis of A549 cells infected with SZ19 (MOI=1) for 24h. Cell lysate was probed with respective nanobodies and detected using goat anti-human IgG-Fc HRP. Data is representative of at least two independent experiments. (F) Sequence of six different nanobodies divided by different area.

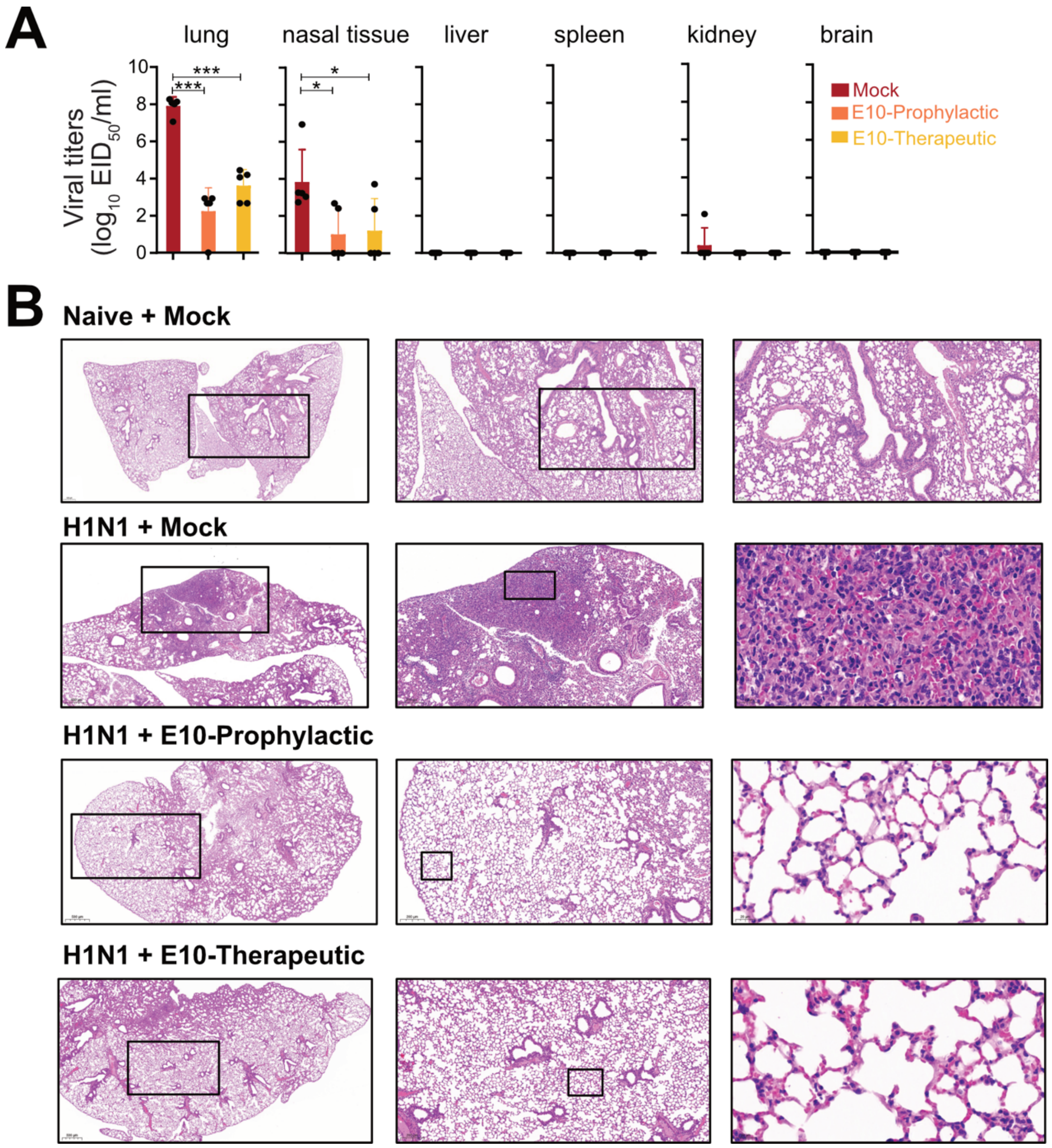


**Figure S2. E10-Fc treatment protects mice against homo- and heterosubtypic IAV challenge.** (A) Viral titer, measured by $EID_{50}$, in six different organs (lung, nasal, liver, spleen, kidney, brain) on day 3 after H1N1 infection with or without E10, as outlined in 2E. Graphs shows mean +/- SD. (B) Representative histopathological analysis of lungs from mice on day 3 after H1N1 infection with or without E10, and with MOCK mice, as outlined 2E. Scar bar, 500 μm in the left row, 200 μm in the middle row, 20 μm in the right row.

A

| Nanobody | Escape mutants (eggs) | Escape mutants (cells) |
| --- | --- | --- |
| A11 | K140N | A122P, K140S |
| C11 | - | - |
| F3 | - | - |
| H10 | S167L, G205E | S167L |
| H12 |  | K140R, G144E, K173E |

B

| Nanobody | Sequences enriched (phage display) | Position |
| --- | --- | --- |
| A11 | VNEEALRQILRESGGIDKEA<br>RESGGIDKEAMGFTYNGIRT | 113-122 |
| C11 | QFELIDNEFNEVEKQIGNVI*3 | 391-410 |
| F3 | AGFIENGWEGLIDGWYGFRH<br>LIDGWYGFRHQNAQGEGTAA | 346-355 |
| H10 | TVTFSFNGAFIAPDRASFLR*3 | 241-260 |
| H12 | IERREGSDVCYPGKFVNEEA<br>YPGKFVNEEALRQILRESGG<br>VNEEALRQILRESGGIDKEA | 103-107 |

C

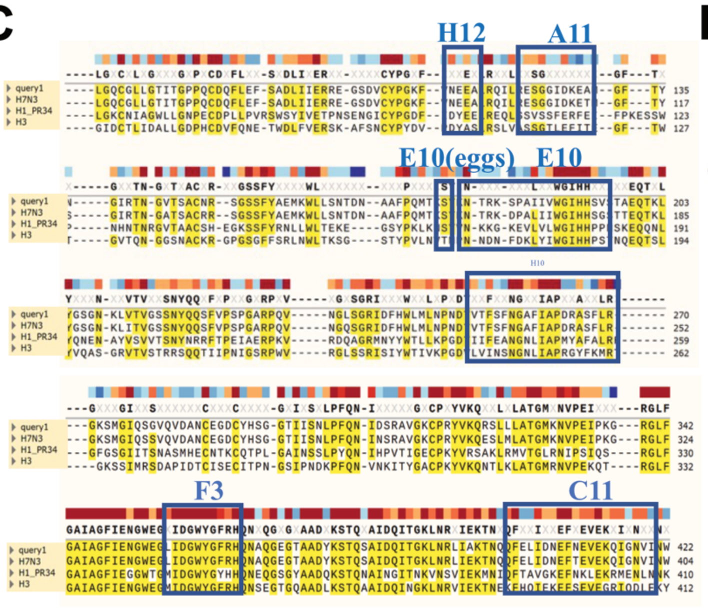


D

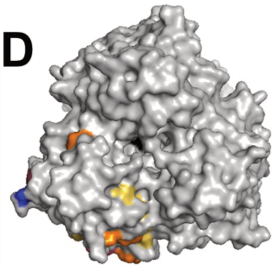

**Figure S3. E10 recognizes a conserved epitope located on HA-head lateral patch.** (A) Nanobody escape mutations in the H7N9 virus were identified using SPF eggs and MDCK cells. Variations at key residues involved in nanobody escape are highlighted, showing the sites where resistance developed. (B) Phage display selection of other 5 nanobody identified specific peptides. The red region depicts an overlapping area among selected peptides. The bolded region corresponds to the epitope identified through escape mutations, which overlap with key binding residues. (C) HA subtype numbering conversion of H7N9-SZ19- HA protein with H1N1-PR8-HA and H3N2-HA. Alignment is working by the website of NIAID Bioinformatics Resource Centers (https://www.bv-brc.org/). The blue sites mean the area of different nanobodies binding in HA protein. (D) Other view of epitope mapping of the H7N9 HA protein was performed, showing the binding sites of six different nanobodies, each labeled in distinct colors (A11: red, E10: orange, H10: yellow, F3: blue, H12: dark blue, C11: black). The head domain of H7 HA was modeled using Swiss-Model. Images were generated using PyMOL software.

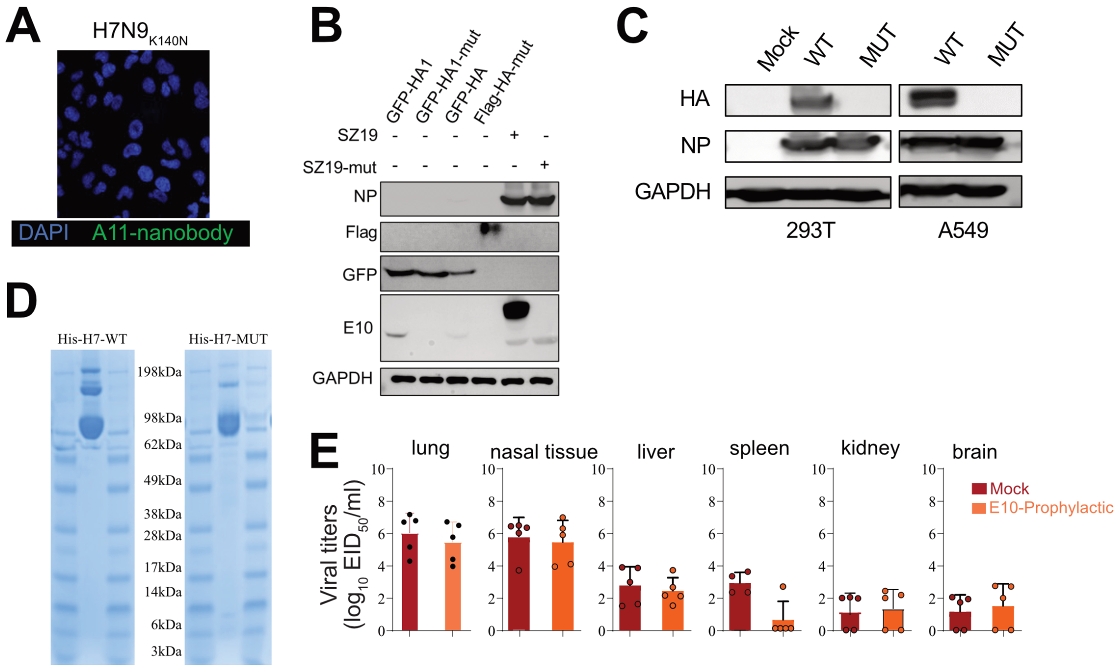


**Figure S4. H7-HA$_{K166T,\ S167L}$ mutants escape E10 recognition but exhibit reduced viral fitness**. (A) IFA of A549 infected with H7N9$_{K140N}$ virus (MOI=0.1) at 24h post-infection. Cell was stained with A11 as the primary antibody, followed by secondary goat anti-human IgG Fc Alexa Fluor™ 488. Nuclei were counterstained with DAPI (blue), and NP nanobody detection is shown in green. Data is representative of at least two independent experiments. (B) Western blot (WB) analysis of A549 cells infected with WT or MUT virus or transfected with GFP-HA and GFP-HA-MUT with or without E10 pre-incubation, showing detection of NP and E10. Data are representative of at least two independent experiments. Shown are the mean values of three technical replicates. (C) Western blot (WB) showing the infection of A549 or 293T cells 24h post infection with WT and MUT virus. Data are representative of at least two independent experiments. Shown are the mean values of three technical replicates. (D) Coomassie Brilliant Blue Staining detect the express and size of recombinant WT-HA and MUT-HA protein. (E) Viral titer, measured by $EID_{50}$, in six organs (lungs, nasal tissue, liver, spleen, kidney, brain) on day 3 post-MUT infection, with or without E10 treatment, as described in 5E. Data are representative of three independent experiments and shown as mean values from three technical replicates. Statistical analysis was performed using unpaired t test. $^*p < 0.05$; $^{**}p < 0.01$; $^{****}p < 0.0001$; ns: not significant.

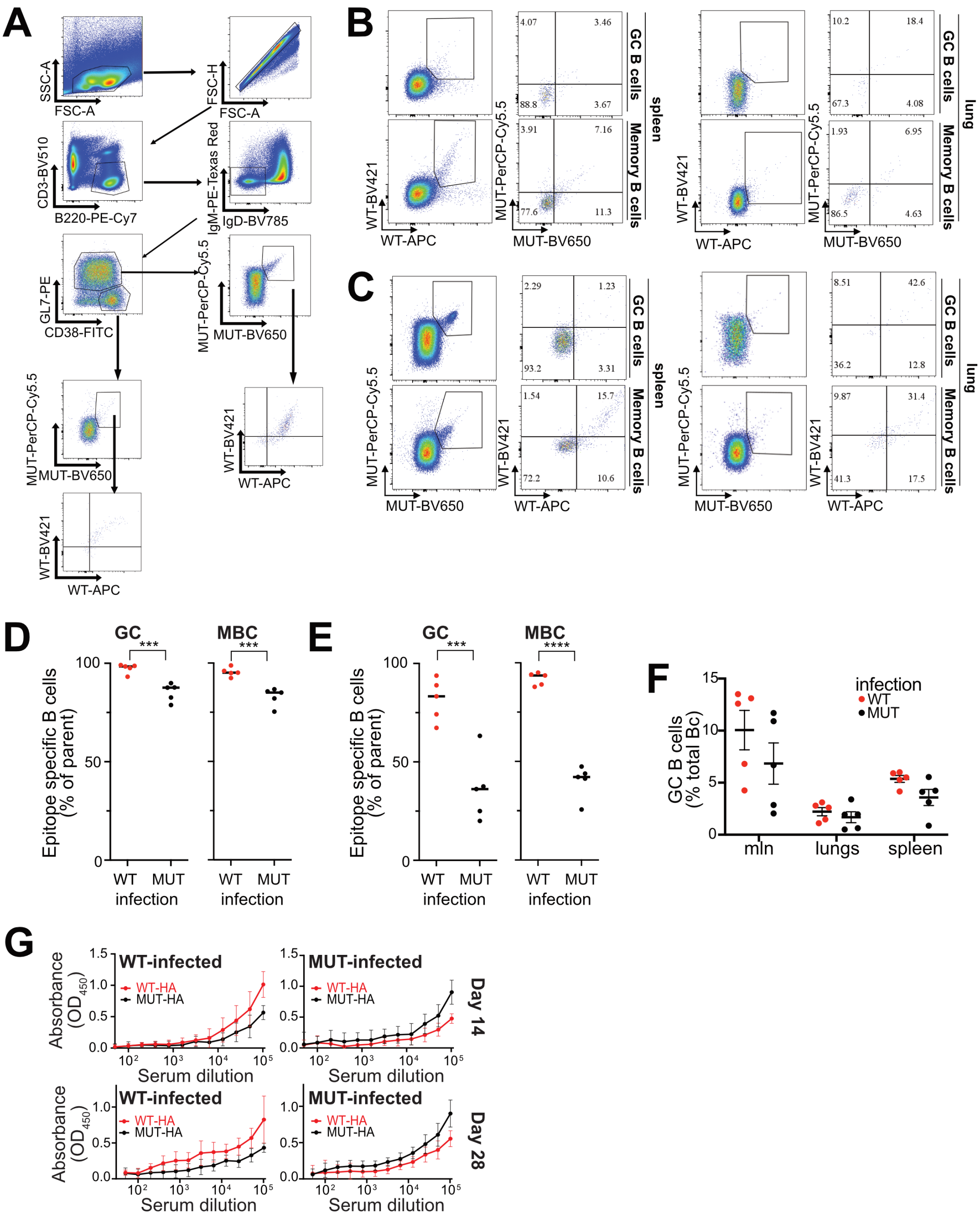


**Figure S5. E10-epitope demonstrates an extreme B cell immunodominance upon H7-IAV infection**. (A) Flow cytometry gating for B cell characterization and analysis. (B) Representative flow cytometry gating of GC and MBC and epitope identification in spleen and lung of 14 days infected WT mice, gated GC as live CD3− B220+ IgD− IgM− GL7+ CD38− WT+ MUT+, MBC as live CD3− B220+ IgD− IgM− GL7− CD38+ WT+ MUT+. (C) Same as in B but for MUT infected mice, gated GC as live CD3− B220+ IgD− IgM− GL7+ CD38− MUT+ WT+, MBC as live CD3− B220+ IgD− IgM− GL7− CD38+ MUT+ WT+. (B, C) Quantification of epitope-specific MBC and GC B cells in lung and spleen of 14 days infected WT and MUT mice. (D, E) Percentage of HA-specific B cells at spleen/lung after infected with WT and MUT virus 14 days. (F) Percentage of B cells at mln/spleen/lung after infected with WT and MUT virus 14 days. (G) ELISA curves showing the detection of WT-HA and MUT-HA in serum of WT and MUT infected mice, at 14, 28 days post infection.

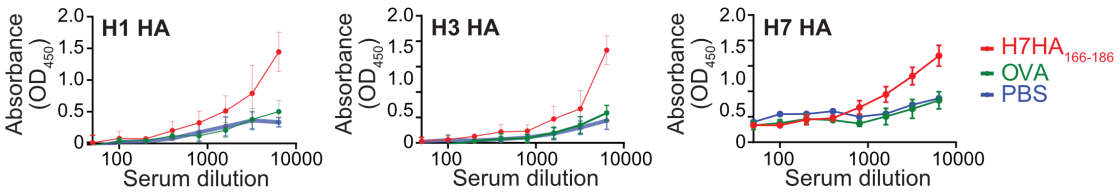


**Figure S6. H7-HA$_{166-186}$ peptide immunization confers partial protection.** ELISA test H1N1/H3N2/H7N9 HA protein can bind with the serum immunized with E10 peptide. (Red line: mice serum which were immunized with H7HA$_{166-186}$ peptide. Green line: mice serum which were immunized with OVA peptide. Blue line: mice serum which were immunized with PBS.)
